# A calibrated cell-based functional assay to aide classification of *MLH1* DNA mismatch repair gene variants

**DOI:** 10.1101/2021.12.14.472580

**Authors:** Abhijit Rath, Alexander A. Radecki, Kaussar Rahman, Rachel B. Gilmore, Jonathan R. Hudson, Matthew Cenci, Sean V. Tavtigian, James P. Grady, Christopher D. Heinen

## Abstract

**PURPOSE:** Functional assays provide important evidence for classifying the disease significance of germline variants in the DNA mismatch repair genes. We sought to develop a cell-based approach for testing the function of variants of uncertain significance (VUS) in the *MLH1* gene.

**METHODS:** Using CRISPR gene editing, we knocked-in *MLH1* VUS into the endogenous *MLH1* loci in human embryonic stem cells. We examined their impact at the RNA and protein level, including their ability to maintain stability of microsatellite sequences and instigate a DNA damage response. We calibrated these assays by testing well-established pathogenic and benign control variants.

**RESULTS:** Five VUS resulted in functionally abnormal protein, 15 VUS resulted in functionally normal protein, and one VUS showed mixed results. Furthermore, we converted the functional outputs into a single odds in favor of pathogenicity score for each VUS.

**CONCLUSION:** Our CRISPR-based functional assay successfully models phenotypes observed in patients in a cellular context. Using this approach, we generated evidence for or against pathogenicity for utilization by variant classification expert panels. Ultimately, this information will assist in proper diagnosis and disease management for suspected Lynch syndrome patients.

## INTRODUCTION

Lynch syndrome (LS; OMIM: 120435) is an autosomal dominant cancer predisposition condition and the predominant cause of inherited colorectal cancer (CRC)^1^. LS patients constitute about 2-4% of all CRC cases^2^. Historically, LS has been associated with increased likelihood of developing colorectal or endometrial cancer. However, recent evidence suggests far greater implications of LS diagnosis in increased predisposition to cancer across multiple tissue types^3^. An LS diagnosis has a profound impact on the clinical management of the patient, but also for instituting proactive cancer screening of “at-risk” relatives. LS diagnosis may be considered due to an early age of cancer presentation, family history, and certain clinical features, although a confirmatory diagnosis of LS depends upon identification of an inactivating germline pathogenic variant in one of the canonical DNA mismatch repair (MMR) genes; *MSH2* (OMIM: 609309), *MLH1* (OMIM: 120436), *MSH6* (OMIM: 600678), and *PMS2* (OMIM: 600259)^1^. Therefore, in recent clinical practice, next-generation sequencing-based identification of MMR gene sequence variants is routinely employed^4^. However, accurate interpretation of the pathogenic significance of gene variants creates a bottleneck, hindering conclusive clinical diagnosis of LS.

To this end, the International Society for Gastrointestinal Hereditary Tumors (InSiGHT), proposed a tiered classification scheme for germline MMR gene variants based on clinical or functional evidence for each variant^5^. Similarly, the American College of Medical Genetics (ACMG) and Association for Molecular Pathology (AMP) generated guidelines for clinical variant interpretation in general based on the same kinds of evidence^6^. For some non-truncating variants, strong family history, variant segregation data, tumor pathology, and population frequency alone is sufficient for classification as benign or pathogenic. However, for nearly 30% of MMR variants identified in the InSiGHT variant database insufficient information exists, resulting in classification as a variant of uncertain significance (VUS)^5,7^. The problem is most evident in the case of a missense change, which constitutes at least 80% of the reported VUS in the four major MMR genes in the ClinVar database.

In the absence of sufficient patient and family history data, laboratory generated functional data can also provide evidence to enable successful classification^8-10^. Thus, the need exists for well-established and validated functional assays. Fortunately, much is known about the normal cellular function of the MMR pathway to provide the basis for such assays. MMR protects genomic integrity by removing a wrongly incorporated nucleotide from the daughter strand during DNA replication^11-14^. MMR also initiates apoptotic signaling in response to persistent DNA adducts, as encountered during treatment of replicating cells with a DNA alkylating agent^15,16^.

Over the course of two decades, multiple laboratories have employed wide-ranging assays for functional interpretation of MMR-gene missense variants^9,10^. However, often a lack of cellular context, use of artificial overexpression systems, or use of non-human model systems prevents a high confidence appraisal of the variants in question.

To address these concerns, we developed a human embryonic stem cell (hESC)-based functional assay using clustered regularly interspaced short palindromic repeats (CRISPR) genome editing to probe MMR gene missense variants^17^. In the current study, we have improved and expanded the use of this approach to assess a large number of *MLH1* VUS, as *MLH1* is the most commonly affected gene in LS patients^7^. We evaluated the ability of these variants to impact MMR-associated repair and damage response functions. In the design of our assay, we followed recommendations from members of the ClinGen Sequence Variant Interpretation Working Group on the use of functional assays in clinical variant interpretation^18^. The strengths of our approach include an *in cellulo* environment which better recapitulates the function of the MMR proteins, the use of genome editing to introduce the variants into the endogenous *MLH1* loci - allowing for simultaneous examination of the effects on RNA and protein, the use of technical and biological replicates to ensure rigor and reproducibility, and the use of known pathogenic and benign variants to allow for assay calibration. By testing a series of established control variants, we were able to use a quantitative framework to convert our functional results into a Bayesian estimate for odds in favor of pathogenicity (OddsPath_Functional) for each of the VUS^19^. This outcome will allow variant interpretation expert panels to readily integrate our data into the existing ACMG/AMP classification scheme.

## MATERIALS AND METHODS

### Generation of *MLH1* variant expressing cell lines

H1 hESCs were obtained from the UConn Stem Cell Core and scored to have a normal karyotype. Cells were cultured on growth factor reduced Matrigel (Corning # 356231) coated plates, fed daily with PeproGrow hESC medium (Peprotech), and maintained at 37°C at 5% CO2 and 95% RH. They were routinely passaged either by microdissection, Relesr (Stemcell Technologies), or StemPro Accutase Cell Dissociation Reagent (ThermoFisher Scientific) upon reaching around 80% confluency. For each *MLH1* (NCBI Gene ID: 4292, RefSeq: NM_000249.4) variant (Table S1), a suitable single guide RNA (sgRNA) to the appropriate genomic target and near a protospacer adjacent motif (PAM) was chosen using an *in silico* program (CRISPOR^20^) (Table S2). Homozygous insertion of the given single nucleotide variant was performed as described previously^17^ with the following modifications: a ribonucleoprotein (RNP)-based targeting approach was used with either a Cas9 or Cas12a recombinant enzyme pre-incubated with the variant-specific Alt-R sgRNAs/crRNAs as per manufacturer’s recommendations (IDT). Following transfection with the Amaxa Stem Cell Nucleofector Kit 2 (Lonza) using an Amaxa Nucleofector II machine, cells underwent selection with 1 μg/mL puromycin (Sigma) for a period of 48 h to isolate single cell clones. Surviving cells were treated with CloneR (STEMCELL Technologies) as recommended. Approximately 32-48 surviving colonies were picked in a PCR hood for each experiment and their genotype confirmed by Sanger sequencing (Table S3 and Fig. S1).

Details of off-target analysis, immunoblotting, exon exclusion assays, cell survival assay, microsatellite instability (MSI) analysis and statistical analyses can be found in the Supplementary Materials and Methods.

## RESULTS

### Generation of *MLH1* variants

We selected 11 known benign and 11 known likely pathogenic/pathogenic control variants and 21 VUS from the InSiGHT database (Fig. **1**). The classification of these variants was performed by the InSiGHT Sequence Variant Interpretation committee, and reassessed without the use of prior functional data in order to avoid circularity in our analyses (Table S1).

**Fig 1.**
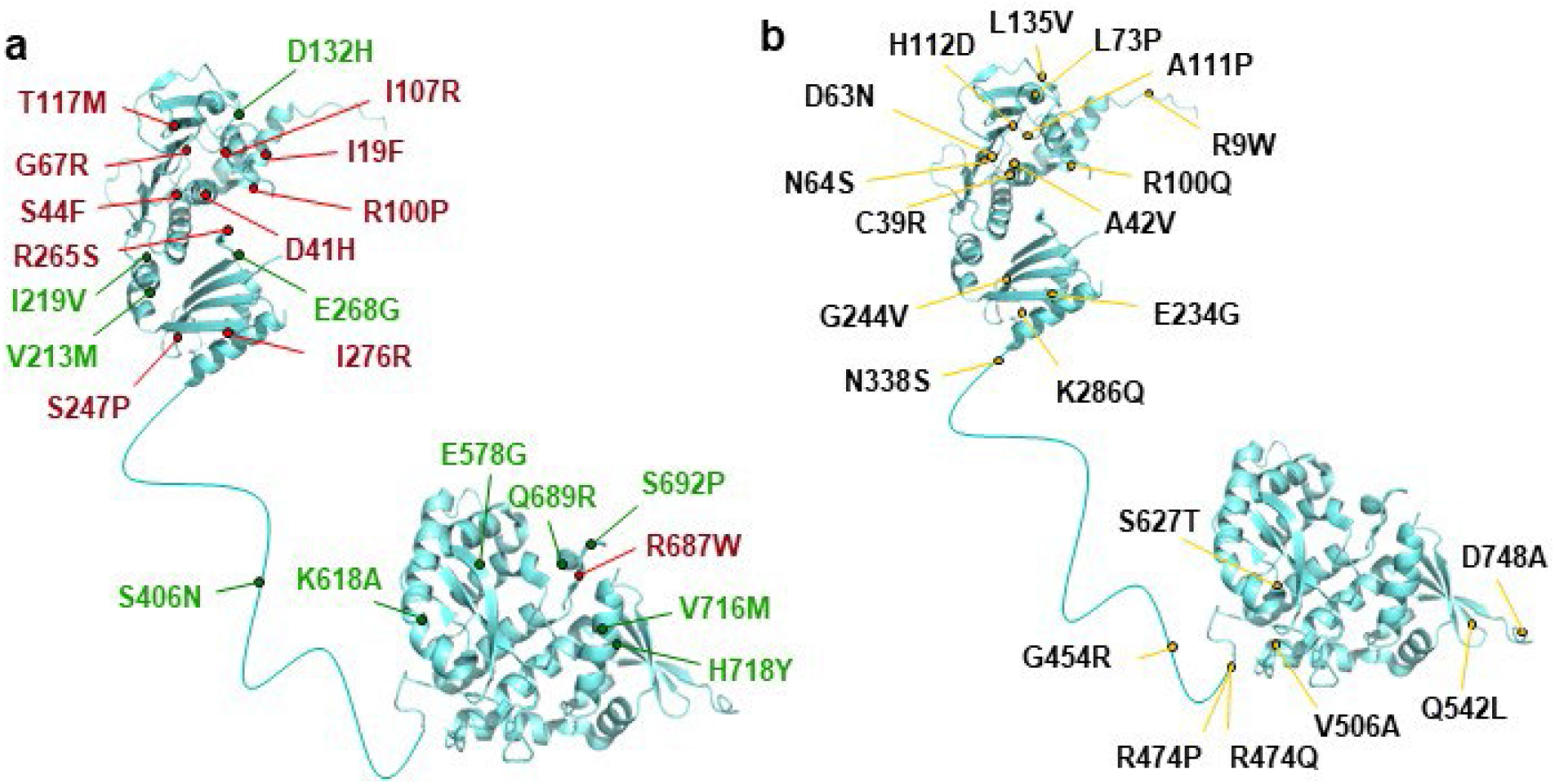
MLH1 variants under study. Representative image of the MLH1 variants in this study mapped to the N-terminal structure of human MLH1 (Protein Data Bank ID code 4P7A) and C-terminal structure of *Saccharomyces cerevisiae* MLH1 (Protein Data Bank ID code 4E4W), connected by a theoretical linker from the human protein. **(a)** The known benign (green) and pathogenic (red) variants selected for functional evaluation. **(b)** The missense variants of uncertain significance (VUS) examined in this study.

We employed CRISPR-Cas gene editing to engineer homozygously targeted cell clones for each variant in wild-type H1 hESCs (WT) (Fig. S1). We also generated an MLH1 knockout cell line (KO) via CRISPR-Cas targeting of exon 2. This targeting resulted in a cell clone in which the MLH1 protein was lost (Fig. **2a**). In general for the variants, we observed a wide-ranging targeting efficiency (Table S4), but the median efficiency was ∼10%, in line with our previous report^17^. Unlike our previous work with *MSH2*, targeting of *MLH1* warranted certain important modifications. First, we opted for a ribonucleoprotein-based approach using commercially sourced sgRNAs, recombinant Cas nucleases, and single strand repair template carrying the single nucleotide variant of interest (Table S2). This approach reduced the reagent preparation time while also drastically reducing the chances of generating unwanted off-target effects^21^. Second, the final 43 cell lines contained only a single nucleotide change to avoid the potential for unintended splicing defects caused by the addition of silent mutations. As a common strategy, introduction of a silent mutation in the PAM site can prevent re-cutting of the sgRNA target sequence by Cas nuclease, and thus increase the chances of recovery of the correctly edited clone. However, in the context of *MLH1*, it may confuse variant interpretation owing to the multiple relatively short exons in *MLH1* and the proclivity for single nucleotide variants of *MLH1* to impart a splicing defect^22^. Instead, we were able to retain a similar editing efficiency by strategically selecting PAM sites, such that the target nucleotide was positioned within the seed sequence of the Cas nuclease target, thereby preventing recurrent cleavage without introduction of an undesired change. Third, use of both Cas9 and Cas12a nucleases offered a greater flexibility to engineer a greater number of variants based on availability of the different PAM sites.

**Fig. 2.**
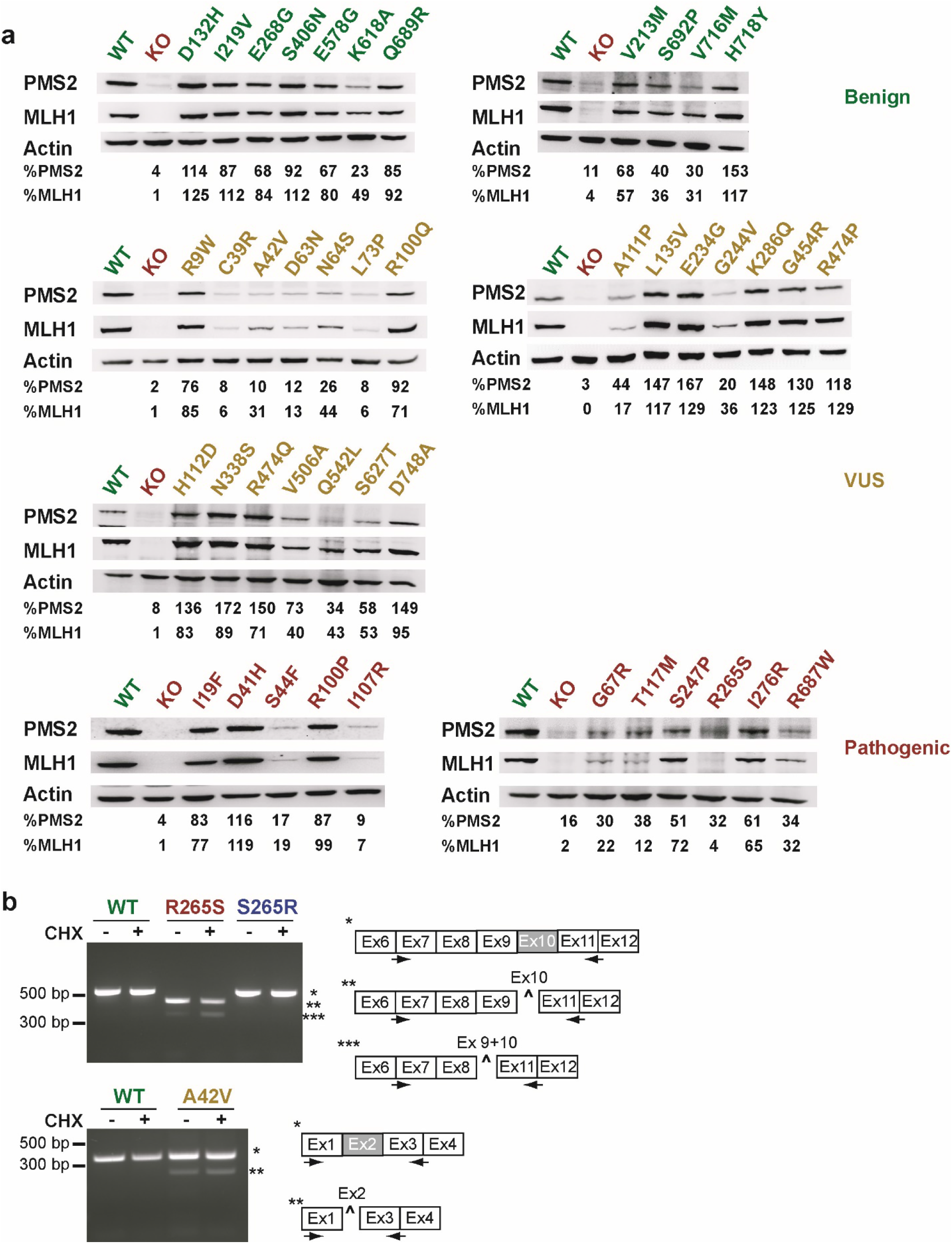
Steady state levels of MLH1 and PMS2 proteins and aberrant splicing variants. **(a**) Representative immunoblots show the steady state levels of MLH1 and PMS2 in benign, pathogenic, and VUS expressing cell lines. Quantification of the protein levels with respect to the WT levels are shown after normalizing for loading. WT, wild-type hESCs; KO, MLH1 knockout hESCs **(b)** Reverse transcribed total RNA from R265S, S265R and A42V cells was PCR amplified to detect the presence of variant containing exons. Representative agarose gel images showing aberrant splicing in indicated cell lines. CHX= Cycloheximide.

To detect evidence of off-target cleavage in the clonally generated cell lines, we bioinformatically identified the most likely off-target sites for each sgRNA. The top five off-target sites were PCR amplified and sequence verified using Sanger sequencing. We did not detect any off-target cleavage in agreement with previous observations from us and others (Table S5)^17,23,24^.

### Impact of missense variants on MLH1-PMS2 steady state level

Loss of stability of MLH1-PMS2 protein product is one way that MMR function is impaired by a pathogenic missense change^9,10^. Thus, as an initial evaluation, we examined steady state levels of MLH1-PMS2 heterodimer in the variant cell lines. As shown in Fig. **2a**., we observed a high degree of variability in MLH1 and PMS2 protein levels both between and within the control variants and VUS compared to WT cells. As expected for known benign controls, MLH1 protein levels remained greater than 80% for most of the 11 variants with the exception of V213M, K618A, S692P, and V716M, which show a greater decrease relative to the WT levels. For the known pathogenic controls, six variants (S44F, G67R, I107R, T117M, R265S, and R687W) showed drastically reduced MLH1-PMS2 levels, while the remaining five (I19F, D41H, R100P, S247P, and I276R) retained relatively stable levels. For the VUS, we observed markedly reduced MLH1 levels (<20% of WT) for C39R, D63N, L73P, and A111P, while the remaining 16 variants showed a varying degree of expression. Since MLH1-PMS2 exists as an obligate heterodimer^25^, an MLH1 variant impacting this interaction can also ultimately affect PMS2 stability. Not surprisingly then, we found that the pattern of PMS2 levels among the variants mostly followed that observed for MLH1.

### Variant induced impact on MLH1 mRNA splicing

One manner in which MLH1 protein levels may be affected by variants is through disruption of mRNA splicing. An advantage of our CRISPR editing-based approach is that we can assess the impact of any variant at both the protein and RNA level. Thus, for each VUS that showed a dramatically reduced protein level (C39R, A42V, D63N, N64S, L73P, A111P, G244V), we examined the impact of the variant on its endogenous mRNA exon. We also examined a set of pathogenic controls that displayed reduced protein levels (S44F, R100P, I107R, T117M, S247P, and R265S). We observed evidence of aberrant splicing for two variants, the pathogenic control R265S located in exon 10 and the VUS A42V located in exon 2. As shown in Fig. **2b**, we failed to detect the expected full-length PCR product spanning exons 7-11 in cDNA from R265S cells, but rather detected two lower migrating bands. Upon gel extraction and Sanger sequencing of the aberrant bands, we confirmed a predominant excision of exon 10 leading to shortening of the amplicon (Fig. S2a). We also detected a fraction of mRNA where both exons 9 and 10 were excluded. Of note, the c.793C>A (R265S) variant has been reported to confer a splicing defect in patient-derived lymphocytes^26^, thus suggesting our CRISPR-based approach can detect variants that impact splicing. To validate the specificity of the variant for this aberrant splicing phenotype, we re-edited the R265S cell line using CRISPR-Cas targeting to re-establish the WT sequence (S265R). We observed a nearly complete rescue of the splicing defect and MLH1 steady-state protein level in the S265R cell line (Fig. **2b** and S3a). The A42V cells, on the other hand, still display a majority of the full-length PCR product spanning exons 1-3 as in WT cells, however, a weaker lower migrating band was observed suggesting exon exclusion (Fig. **2b**). Sanger sequencing of the aberrant splice product confirmed exclusion of the exon 2 (Fig. S2b). The remaining variants tested all displayed normal exon splicing, suggesting that the changes in protein levels are likely due to impacts of the variant on protein stability (Fig. S4 and Table S6).

### Variant effect on DNA damage signaling

Mammalian cells mount a cell cycle arrest and/or apoptotic response upon treatment with a DNA alkylating agent in an MMR-dependent fashion^15,16^. To assess the ability of *MLH1* VUS expressing cells to successfully instigate such a response, we measured cell survival 48 h post treatment with the S_N_1 alkylating agent MNNG. We also subjected WT and MLH1 KO cells as well as the known benign and pathogenic controls to MNNG. As expected, greater than 80% of MLH1 KO cells survived MNNG treatment, whereas only 15% of WT cells did (Fig **3**). The eleven benign controls responded similarly to the WT cells, though two cell lines, E578G and K618A, displayed slightly increased resistance to MNNG. The known pathogenic variant expressing cells demonstrated resistance to treatment similar to KO cells. To evaluate the VUS, we used statistical clustering to identify those variants whose survival level was similar to either benign or pathogenic controls. Two, non-overlapping clusters were generated demonstrating a clean separation of the results for the benign and pathogenic controls. Of the 21 MLH1 VUS examined, C39R, D63N, L73P, R100Q, A111P, and G244V showed resistance to MNNG, clustering with the pathogenic controls. The remaining 16 VUS clustered with the benign controls. Two variant lines, A42V and S627T, demonstrated a slightly increased resistance to MNNG compared to WT cells, though still segregated with the benign cluster. As an additional control to confirm that the change in damage response in a pathogenic variant line was indeed due to the sequence variant, we tested the response to MNNG in the S265R revertant cell line. We found that re-editing this variant back to the WT sequence restored the normal response to DNA damage (Fig. S3b).

**Fig. 3.**
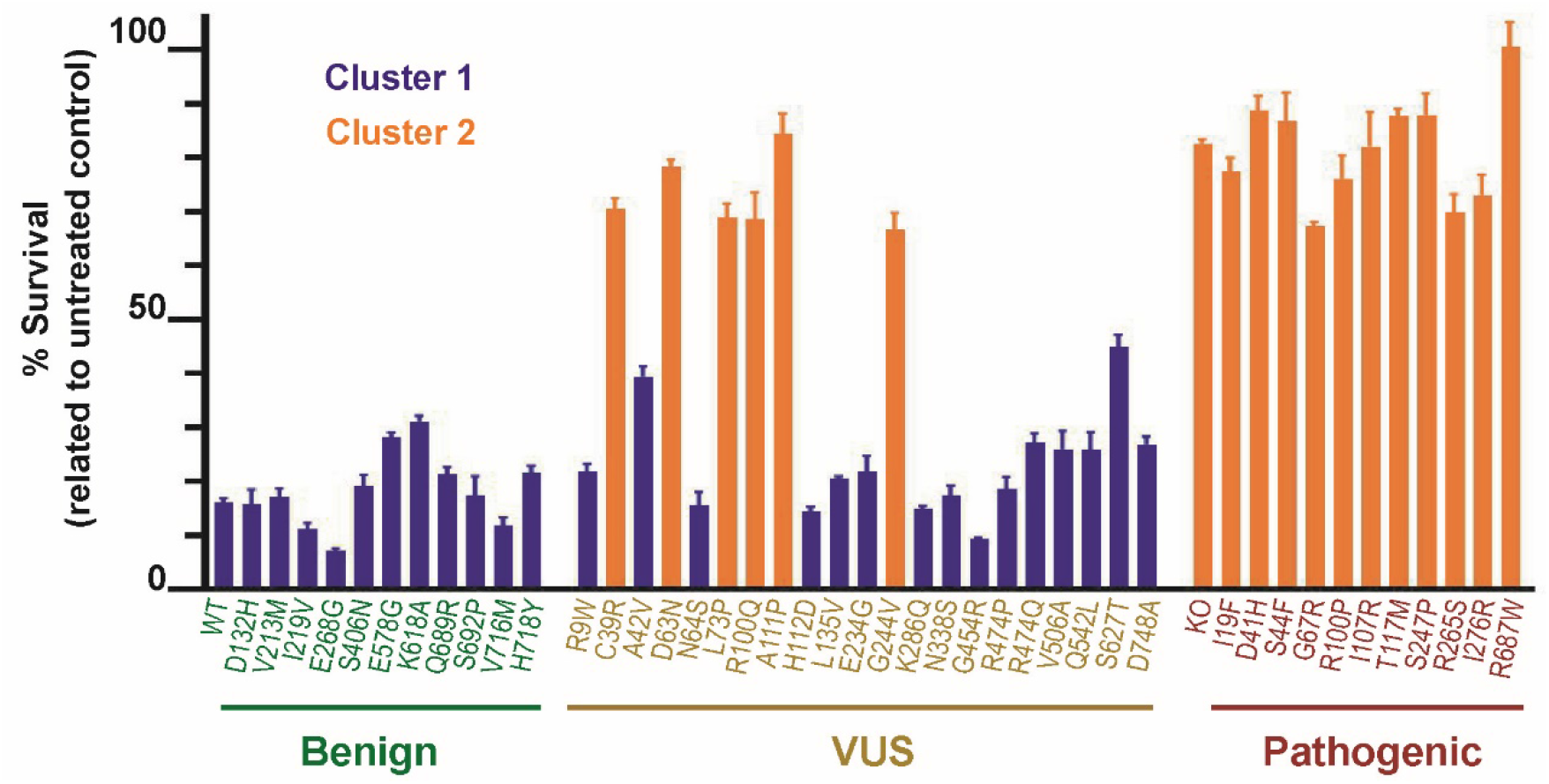
The mismatch repair-dependent damage response in variant cell lines. Cell survival after 48 hr treatment with 2 μM of the DNA alkylating agent *N*-methyl-*N’*-nitro-*N*-nitrosoguanidine. The values are represented as the mean ± SEM; n=3-5. A statistical clustering analysis was performed which grouped the variants into two populations based on their percent survival (blue and orange).

### Examining microsatellite instability (MSI) in *MLH1* VUS

Deficient MMR function leads to instability in microsatellite repeats (MSI) and the presence of MSI is the best-characterized molecular evidence for LS associated tumors^27^. We examined MSI in the *MLH1* VUS expressing cells as another read-out for MMR function. The benign and pathogenic variants were also examined to serve as controls. We passaged each cell line for 15-20 generations and then isolated 32 single cell clones to evaluate for MSI at five mononucleotide repeats (Table 1 and Table S7). We observed few or no unstable clones in the WT and benign control cells, with higher numbers of unstable clones in the KO and pathogenic controls. To more quantitatively determine if MSI can be used to distinguish benign and pathogenic variants, we again employed a statistical clustering approach. Using the results for each cell line from all five MSI markers, we asked whether the benign and pathogenic controls would segregate into two separate clusters. We observed this distinct clustering of controls for three markers, BAT-25, BAT-26 and NR-27 with only two discordant results (G67R for BAT-26 and NR-27) (Fig. S5). However, the controls failed to segregate into two separate clusters for the NR-21 and NR-22 markers (Fig. S5), suggesting that these MSI markers may not be informative for distinguishing variants.

**Table 1.** Microsatellite instability in MLH1 variant-expressing cell lines.

| **Variant/MSI Marker** | **NR-21<sup>a</sup>** | **NR-22<sup>a</sup>** | **NR-27<sup>a</sup>** | **BAT-25<sup>a</sup>** | **BAT-26<sup>a</sup>** |
| --- | --- | --- | --- | --- | --- |
| WT | 0/30 | 0/31 | 0/29 | 0/30 | 0/31 |
| KO | 28/32 | 30/30 | 31/31 | 15/31 | 31/31 |
| *Benign* |  |  |  |  |  |
| D132H | 0/30 | 0/31 | 0/32 | 0/32 | 0/32 |
| V213M | 1/32 | 0/32 | 0/30 | 1/31 | 0/32 |
| I219V | 5/32 | 0/32 | 0/30 | 0/32 | 1/32 |
| E268G | 1/32 | 0/32 | 0/32 | 0/32 | 0/31 |
| S406N | 6/32 | 0/32 | 0/28 | 0/29 | 0/32 |
| E578G | 2/32 | 0/32 | 1/27 | 1/28 | 0/32 |
| K618A | 0/32 | 0/32 | 0/32 | 1/32 | 0/32 |
| Q689R | 5/30 | 0/30 | 0/29 | 0/29 | 0/31 |
| S692P | 7/32 | 0/31 | 6/30 | 0/29 | 2/31 |
| V716M | 0/31 | 0/31 | 1/31 | 0/31 | 0/30 |
| H718Y | 4/32 | 0/32 | 0/32 | 0/32 | 4/32 |
| *VUS* |  |  |  |  |  |
| R9W | 6/32 | 0/32 | 11/32 | 0/31 | 2/32 |
| C39R | 25/32 | 14/32 | 32/32 | 17/32 | 21/32 |
| A42V | 2/32 | 1/32 | 0/32 | 0/32 | 0/32 |
| D63N | 13/30 | 27/29 | 13/30 | 23/30 | 28/31 |
| N64S | 0/32 | 0/32 | 0/32 | 0/32 | 0/32 |
| L73P | 24/31 | 5/31 | 22/31 | 21/29 | 16/32 |
| R100Q | 13/32 | 2/32 | 9/31 | 9/32 | 6/32 |
| A111P | 15/32 | 14/32 | 15/32 | 16/32 | 20/32 |
| H112D | 0/32 | 5/32 | 0/31 | 0/32 | 0/31 |
| L135V | 2/32 | 0/32 | 0/32 | 1/32 | 0/32 |
| E234G | 3/32 | 0/32 | 3/31 | 0/32 | 0/32 |
| G244V | 25/28 | 6/28 | 15/26 | 15/26 | 18/27 |
| K286Q | 8/31 | 0/31 | 4/28 | 0/31 | 0/31 |
| N338S | 0/32 | 1/32 | 0/32 | 0/32 | 0/32 |
| G454R | 0/32 | 0/32 | 0/32 | 0/32 | 0/32 |
| R474P | 0/32 | 0/32 | 0/31 | 0/32 | 0/32 |
| R474Q | 6/31 | 0/30 | 0/29 | 0/30 | 0/31 |
| V506A | 0/31 | 0/29 | 0/25 | 0/29 | 0/30 |
| Q542L | 0/27 | 0/31 | 0/31 | 0/31 | 0/32 |
| S627T | 10/29 | 0/29 | 0/29 | 0/29 | 0/32 |
| D748A | 1/32 | 0/32 | 0/30 | 0/32 | 0/32 |
| *Pathogenic* |  |  |  |  |  |
| I19F | 21/32 | 11/32 | 23/32 | 13/32 | 30/32 |
| D41H | 13/32 | 21/31 | 15/30 | 16/31 | 23/32 |
| S44F | 15/31 | 10/31 | 29/31 | 30/31 | 28/31 |
| G67R | 27/32 | 7/32 | 13/31 | 16/30 | 10/31 |
| R100P | 15/31 | 5/31 | 22/32 | 24/32 | 18/32 |
| I107R | 15/32 | 18/29 | 17/28 | 12/25 | 25/31 |
| T117M | 14/29 | 11/29 | 22/28 | 17/25 | 22/29 |
| S247P | 31/32 | 10/32 | 17/29 | 11/32 | 23/32 |
| R265S | 15/32 | 7/31 | 28/32 | 23/28 | 29/30 |
| I276R | 17/32 | 13/31 | 25/25 | 17/30 | 19/31 |
| R687W | 14/30 | 6/30 | 23/28 | 11/27 | 25/31 |

<sup>a</sup> Ratio of clones with altered alleles compared with WT sequence to total clones tested

### Odds of Pathogenicity Scores

In order to convert the results from our functional assays into a value that can be utilized in the ACMG/AMP classification scheme, we utilized a Bayesian framework that allows for the expression of evidence into a quantitative OddsPath score^19^. To generate this score, we obtained previously established OddsPath values for the 11 pathogenic and 11 benign variants that were derived based on existing personal and clinical data (Table S1). The contribution of any prior functional assay was removed from the calculations in order to avoid circularity^18^. To calibrate our functional assays, linear regression analyses were performed with the OddsPath score as the dependent variable and the assay readout as the independent variable. We initially obtained a single OddsPath score from the MSI assay for each VUS by averaging the results from the three informative MSI markers (BAT25, BAT26 and NR27). A separate OddsPath score was derived from the MNNG survival data and then both functional scores were combined to create a single OddsPath_Functional score for each VUS (Table 2). Using the draft ACMG/AMP classification criteria for MMR genes (https://www.insight-group.org/criteria/), we assigned the appropriate evidence code corresponding to these OddsPath_Functional scores. Five variants had OddsPath_Functional scores > 18.7 consistent with strong evidence of Pathogenicity (PS3), whereas 15 VUS had OddsPath_Functional scores < 0.05, consistent with strong evidence of a benign classification (BS3). We then combined these new evidence codes with existing clinical, genetic and *in sillico* evidence available for each variant to determine a predicted classification. From these results, we expect that 13 VUS will be re-classified (Table 2). One variant that remains a VUS is R100Q, which had an OddsPath_Functional score of 0.778, which is indeterminate for pathogenicity.

**Table 2.** Summary of Functional Study Outcomes and Predicted Classifications for VUS.

| Variant | Protein Stability<sup>a</sup> | Damage Response Signaling<sup>a</sup> | DNA Repair Function<sup>a</sup> | OddsPath_Functional Score<sup>b</sup> | ACMG/AMP Evidence Code<sup>c</sup> | Predicted Classification<sup>d</sup> |
| --- | --- | --- | --- | --- | --- | --- |
| R9W | + | + | + | 8.70E-5 | BS3 | VUS |
| C39R | - | - | - | 508 | PS3 | LP |
| A42V | +/- | + | + | 2.54E-4 | BS3 | VUS |
| D63N | - | - | - | 1,350 | PS3 | LP |
| N64S | +/- | + | + | 5.65E-6 | BS3 | B |
| L73P | - | - | - | 108 | PS3 | LP |
| R100Q | + | - | +/- | 0.778 | Ind | VUS |
| A111P | - | - | - | 379 | PS3 | LP |
| H112D | + | + | + | 4.58E-6 | BS3 | VUS |
| L135V | + | + | + | 1.31E-5 | BS3 | LB |
| E234G | + | + | + | 2.19E-5 | BS3 | LB |
| G244V | +/- | - | - | 57.5 | PS3 | LP |
| K286Q | + | + | + | 9.15E-6 | BS3 | VUS |
| N338S | + | + | + | 7.25E-6 | BS3 | LB |
| G454R | + | + | + | 1.95E-6 | BS3 | LB |
| R474P | + | + | + | 8.63E-6 | BS3 | LB |
| R474Q | + | + | + | 3.11E-5 | BS3 | LB |
| V506A | +/- | + | + | 2.60E-5 | BS3 | VUS |
| Q542L | +/- | + | + | 2.55E-6 | BS3 | VUS |
| S627T | + | + | + | 6.41E-4 | BS3 | LB |
| D748A | + | + | + | 2.97E-5 | BS3 | VUS |

<sup>a</sup> From this study
<sup>b</sup> OddsPath\_Functional scores calculated as described in the Methods using the known benign and pathogenic variants for calibration
<sup>c</sup> ACMG/AMP evidence code derived from OddsPath scores in accordance with the draft ACMG/AMP classification criteria for MMR genes (https://www.insight-group.org/criteria/). PS3, strong pathogenic; BS3, strong benign; Ind, indeterminate
<sup>d</sup> Predicted ACMG/AMP classification derived from combining OddsPath\_Functional score with prior *in silico*, clinical or genetic data (see Table S1) using draft ACMG/AMP classification criteria for MMR genes. LP, likely pathogenic; LB, likely benign; B, benign

## DISCUSSION

The ACMG/AMP guidelines for clinical variant interpretation include the use of functional studies as one means to predict the impact of a genetic variant^6^. A point of emphasis in these guidelines is that researchers should consider the degree to which a functional assay reflects the biological environment and disease mechanism. The CRISPR-based approach to study MMR gene variants described here examines each variant in the context of the human cell, which allowed us to determine the impact of variant proteins in their normal nuclear environment. The use of a cell-based assay also allowed us to examine both repair of endogenous microsatellites and the initiation of a global cell response to DNA damage, both mechanisms that are likely involved in disease etiology^1,15,28^. By introducing the variant into the endogenous *MLH1* gene loci, we ensured that the *MLH1* endogenous promoter and regulatory sequences control its expression. This also allowed us to assess the effects of the variant on the *MLH1* RNA and protein simultaneously. By doing so, we found that two variants altered mRNA splicing, R265S and A42V. Whereas the R265S variant led to exclusion of exon 10, or 9 and 10, in all transcripts, resulting in an abnormally functioning protein, the A42V variant had only a partial defect, producing enough full-length protein to retain normal function. Combining mRNA analyses with MMR pathway functional testing in the same cells allows us to resolve challenges that arise in interpreting bioinformatic predictions or *ex vivo* assays of mRNA splicing^29^.

To align the output of functional assays such as ours with the ACMG/AMP guidelines, one of us previously derived a quantitative approach to convert a functional result into an OddsPath score that could then be converted to ACMG/AMP evidence codes for or against pathogenicity^19^. Adopting this overall approach, members of InSiGHT created draft ACMG/AMP guidelines specific to *MSH2, MSH6, MLH1* and *PMS2* (https://www.insight-group.org/criteria/). In these guidelines, OddsPath scores > 18.7 indicate strong evidence for pathogenicity (PS3). Based on our data, five VUS can be assigned this evidence strength (C39R, D63N, L73P, A111P and G244V; see Table 2). The remainder of the VUS tested, with one exception, all had OddsPath scores < 0.05, indicative of strong evidence for a benign classification (BS3). However, functional results are not considered to be stand-alone evidence for a pathogenic or benign classification^18^. Thus, functional evidence must be combined with at least one other piece of data including *in silico*, clinical or genetic data for the given variant. Using prior data available on the InSiGHT database for the 21 VUS tested here, we generated classification predictions (Table 2). Five variants are expected to reach the level of likely pathogenic, while 7 variants are expected to reach the level of likely benign. One variant (N64S) reached the level of benign due to another existing piece of strong benign evidence. This variant was reported to co-occur with a known pathogenic *MLH1* variant in a patient who did not demonstrate a Constitutive MMR-Deficiency Disease phenotype (evidence code BS4)^30^. The 8 remaining variants should still be considered VUS at this time, most of them due to discordance between our functional results and prior *in silico* evidence from MAPP and PolyPhen-2. We have observed some discordance between *in silico* data and functional data in our previous study of *MSH2* variants as well^17^, as the *in silico* approaches tend to over-predict pathogenicity. This result further highlights why neither approach should be used alone for direct classification.

The one OddsPath_Functional score exception was for R100Q, which had a score of 0.778, indicating indeterminate evidence for classification. Interestingly the R100Q cells displayed a lack of response to DNA damage similar to a functionally abnormal variant, but only limited MSI. This contrasts with all other functionally abnormal variants, which showed both the loss of damage response and high levels of MSI. Of note, an analogous change in the corresponding amino acid for *Escherichia coli* MutL (R95F) has undergone interrogation in single molecule studies^31^. Because of its location in the hinge region of the ATP lid of the MutL/MLH1 protein, the R95F variant inhibits ATP binding. While this variant does not prevent ternary complex formation between MutL, MutS, and DNA, it does impact the formation of MutL sliding clamps on DNA. Future studies will be needed to determine whether the R100Q variant similarly reduces the levels of MLH1-PMS2 sliding clamps on DNA. Why this variant appears so disruptive for the DNA damage signaling function of MLH1, but not quite as disruptive for repair of microsatellite sequences is unclear. One hypothesis is that the damage signaling function requires more efficient MLH1-PMS2 sliding clamp formation compared to standard microsatellite repair. This would be consistent with previous observations that the damage signaling function is more sensitive to significantly reduced levels of MLH1 expression than standard repair ^32^. However, the effect of MLH1-PMS2 protein levels is not quite as clear from our data. A number of variants appear to impact protein levels that then result in reduced MMR function, however, some variants show significantly reduced protein levels, yet maintain normal MMR function (Table 2). This includes a Class 1 benign variant (K618A) along with several VUS that have OddsPath_Functional scores consistent with benign interpretation. Interestingly, of those VUS with reduced protein levels, all but one show slightly poorer DNA damage responses than the wild-type protein including two variants that are borderline intermediate in their response (A42V and S627T; Fig. **3**). Yet, all of these variants have very stable microsatellites. Overall, these results suggest that the DNA damage response may be more sensitive to the levels of MLH1-PMS2 sliding clamp formation that is different from regular repair. The potential for a mechanistic separation between repair and the damage response, particularly as seen with the R100Q variant, will be interesting to examine in future studies to help clarify the molecular mechanism of the MMR-dependent damage signaling response. In contrast to R100Q, another variant tested here, G67R, had previously been reported as a separation of function allele when engineered in mice^33^. *Mlh1*^*G67R/G67R*^ mouse embryonic fibroblasts retained an apoptotic response to the DNA crosslinking agent cisplatin, even though the *Mlh1*^*G67R/G67R*^ animals developed tumors marked by MSI similar to *Mlh1*^*-/-*^ mice. However, in our studies, we found that the human G67R variant led to both increased MSI and a loss of DNA damage sensitivity. While species-specific differences are one possible explanation, it may also speak to differences in the mechanism of MMR-dependent signaling generated by crosslinking damage compared to DNA alkylation.

In summary, our CRISPR based functional assays provide a means to assess the effects of MMR gene VUS in a human cellular context. By calibrating the assays with multiple known pathogenic and benign controls, an OddsPath_Functional score can be derived for direct use in variant classification schemes. While we have provided predictions for how these results may impact classification, ultimately, this task will require review by an expert panel such as the InSiGHT variant interpretation committee^5^. The challenge that remains is to perform these functional assays in a more high-throughput manner in order to more rapidly address the many remaining MMR gene VUS. A recent high-throughput approach examining *MSH2* variants was reported, providing encouragement that this large-scale problem may be manageable in the near future^34^. However, in this study, the researchers used lentivirus to express variant-containing *MSH2* cDNAs. This approach has a level of artificiality with regards to regulation of gene expression, while also missing a potential role for variants in splicing anomalies. Their assay also did not examine direct repair function such as maintaining stability of microsatellites. Thus, confidence in using these results for classification may be limited. The ability to scale-up the use of CRISPR gene editing for testing VUS may help overcome these challenges.

## Data Availability

Information on variants tested in this study can be found at https://www.insight-database.org/classifications/mmr_integrative_eval.html. We will supply all other data and materials upon request.

## Acknowledgments

The authors wish to thank John-Paul Plazzer for his assistance in obtaining relevant variant data from the InSiGHT database and advising on the use of the MMR gene draft classification criteria. This work was supported by the National Institutes of Health grants CA222477 and CA237920 (C.D.H.), and CA164944 (S.V.T.).

## Author Information

Conceptualization: Ab.R., S.V.T., J.P.G., C.D.H; Data curation: Ab.R., A.A.R., J.P.G., C.D.H.; Formal Analysis: Ab.R., J.P.G., C.D.H.; Funding Acquisition: S.V.T., J.P.G., C.D.H.; Investigation: Ab.R., A.A.R., K.R., R.B.G., J.R.H. M.C., J.P.G.; Methodology: Ab.R., S.V.T., J.P.G., C.D.H.; Project Administration: C.D.H.; Supervision: C.D.H.; Validation: Ab.R., A.A.R., K.R.,C.D.H.; Writing – original draft: Ab.R., C.D.H.; Writing – reviewing & editing: Ab.R., A.A.R., K.R., R.B.G., J.R.H., M.C., S.V.T., J.P.G., C.D.H.

