## Supplementary Materials for "A calibrated cell-based functional assay to aide classification of *MLH1* DNA mismatch repair gene variants"

### SUPPLEMENTARY MATERIALS AND METHODS:

#### Off-target analysis

A list of probable top five off-target binding sites for each sgRNA used in this study was determined using the CRISPOR website[1]. CRISPOR version 4.97 was used to obtain the off-target loci according to the Cas nuclease used in each targeting. Each site was PCR amplified using high fidelity Phusion DNA polymerase (NEB) and analyzed by Sanger sequencing (Table S5). The integrity of each site was verified by carrying out pairwise sequence alignment with the reference genome along with manual examination of the DNA chromatogram.

#### Immunoblotting

Approximately one million cells for each cell line were lysed in 120 µL RIPA buffer in the presence of protease inhibitors. 25 µg of each cell lysate was run on an 8% SDS-PAGE gel and probed with anti-MLH1 (BD Pharmingen # 554073, 1:1000), anti-PMS2 (BD Pharmingen # 556415, 1:1000), and anti-actin (Santa Cruz Biotechnology # sc-8432, 1:1000) antibodies. Quantitation of MLH1 and PMS2 levels in variant expressing lines was carried out in comparison to MLH1 or PMS2 levels in WT cells after being normalized for protein loading using actin levels via ImageJ software.

#### Exon inclusion assay

RNA splicing analyses were carried out as described previously[2,3]. Briefly, indicated cell lines were treated with 20 µg/mL cycloheximide (Sigma-Aldrich) for 4 h to suppress nonsense-mediated decay and allow accumulation of potential unstable transcripts. Subsequently, 500 ng of total RNA was used in a reverse transcription reaction using the iScript cDNA synthesis kit (Bio-Rad). Each variant-containing exon was detected by PCR amplification using primers in the flanking exons (Table S6). The PCR products were analyzed on a 1.5% agarose gel.

**MNNG drug toxicity assay**

Approximately 4,000 cells of each cell line were seeded into individual wells of a Matrigel coated 48-well plate along with 10 μM ROCK inhibitor (Y-27632, Selleck Chemicals). The medium was replaced without the ROCK inhibitor the next day and cells were allowed to grow overnight. Approximately 36 h after seeding, cells were initially treated with 25 μM $O^6$-Benzylguanine ($O^6$-BG; Sigma) for 2 h to inhibit action of the $O^6$-methylguanine-DNA methyltransferase (MGMT). Subsequently, cells were treated with 2 μM of *N*-methyl-*N'*-nitro-*N*-nitrosoguanidine (MNNG) (obtained from the National Cancer Institute Chemical Carcinogen Reference Standard Repository) for 48 h in the presence of $O^6$-BG and surviving cell population was assessed using the 3-(4,5-dimethylthiazol-2-yl)-2,5-diphenyltetrazolium bromide (MTT) assay (ThermoFisher Scientific) following the manufacturer's instruction. Assays were performed in triplicate and repeated at least 3 times.

**Microsatellite instability (MSI) analysis**

For each cell line, the cell population was extensively diluted to isolate and grow multiple single cell colonies for 7–10 days. Approximately 32 single cell clones were picked for each line and genomic DNA was extracted. All the cell lines used for MSI analysis were between passage number 15-20. Five mononucleotide loci NR-21, NR-22, NR-27, BAT-25 and BAT-26 were amplified using standard Taq DNA polymerase (NEB) with either 6-FAM or HEX labeled fluorescent forward primers. Two multiplex reactions (NR-27 and BAT-25, NR-21 and NR-22) and a single reaction (BAT-26) were set up for each clone. All three individual PCR products were combined for fragment analysis by capillary electrophoresis (Genewiz). Clones showing changes in fragment length compared to WT genomic DNA were scored as MSI-positive and the percentage of positive clones were calculated for each marker per cell line. Manual scanning of individual peaks along with the Geneious software program were used to identify instability.

#### Statistical analyses

To assess how the VUS expressing cell lines behaved in functional assays relative to the known pathogenic and benign controls, hierarchical clustering was used specifying a two-cluster solution in the CLUSTER procedure in SAS. All control variants and VUS were analyzed simultaneously and the two best clusters identified. This process was performed independently using the percentage survival score for the MNNG survival assay as well as the fraction of unstable clones for each of the five MSI markers. To convert our functional results into an OddsPath_ Functional score, we first obtained previously calculated OddsPath_Non-Functional scores for the known pathogenic and benign control variants based on available clinical and personal data from patients carrying these variants as found at http://insight-database.org/classifications/ (Table S1). These scores were used to estimate a linear relationship on the log10 scale. The log10 OddsPath_Non-Functional scores were treated as the dependent variable and the assay results as the independent variable in a linear regression. The resulting predictive equations were then applied to the VUS to produce log10 OddsPath_Functional score estimates similar to that performed previously [4,5]. The GENMOD procedure in SAS was used to estimate the linear model parameter estimates and their standard errors and the predicted log10 OddsPath_Functional scores were generated using the PSL procedure in SAS. This analysis was performed independently for the MNNG survival assay as well as the MSI assay to generate a log10 OddsPath score for each assay. To generate a single MSI assay score, the fraction of unstable clones from each of the three informative MSI markers (Fig S5) were averaged for each VUS and the predictive equation applied to this mean. A final OddsPath_Functional score was created by combining the MNNG survival assay score and MSI assay score using a meta analytic approach with the GENMOD procedure in SAS, which gives a variance weighted estimate with 95% confidence limits [6].

**Supplementary Table 1 *MLH1* variant characteristics**

| Cell Line | Variant in cDNA | ClinVar ID | OddsPath_Non-functional[a] | Polyphen-2[b] | MAPP-MMR[c] | PON-MMR2[d] |
|---|---|---|---|---|---|---|
| *Benign* | | | | | | |
| D132H | c.394G>C | 17096 | 1.08E-7 | 0.999 Probably damaging | 4.73 | Neutral |
| V213M | c.637G>A | 41640 | 5.76E-4 | 0.107 Benign | 1.99 | Neutral |
| I219V | c.655A>G | 45219 | - | 0.060 Benign | 3.51 | Neutral |
| E268G | c.803A>G | 90382 | 2.24E-4 | 0.979 Probably damaging | 5.20 | Neutral |
| S406N | c.1217G>A | 41634 | 1.49E-10 | 0.081 Benign | 1.07 | Neutral |
| E578G | c.1733A>G | 17090 | 1.44E-6 | 0.041 Benign | 4.85 | Neutral |
| K618A | c.1852_1853delinsGC | 32128 | 2.90E-13 | 0.991 Probably damaging | 5.06 | Neutral |
| Q689R | c.2066A>G | 90016 | 1.11E-4 | 0.001 Benign | 3.36 | Neutral |
| S692P | c.2074T>C | 127621 | 4.02E-4 | 0.000 Benign | 4.64 | Neutral |
| V716M | c.2146G>A | 41639 | 4.07E-5 | 0.704 Possibly damaging | 2.78 | Neutral |
| H718Y | c.2152C>T | 36547 | - | 0.702 Possibly damaging | 3.45 | Neutral |
| *VUS* | | | | | | |
| R9W | c.25C>T | 90122 | - | 0.973 Probably damaging | 4.48 | Pathogenic |
| C39R | c.115T>C | 89655 | 71.4 | 0.949 Probably damaging | 37.7 | Pathogenic |
| A42V | c.125C>T | 89688 | - | 1.0 Probably damaging | 30.5 | Pathogenic |
| D63N | c.187G>A | 89920 | 0.00 | 1.0 Probably damaging | 13.8 | Neutral |
| N64S | c.191A>G | 89947 | 5.00E-2 | 0.994 Probably damaging | 11.8 | Pathogenic |
| L73P | c.218T>C | 90086 | 13.7 | 0.996 Probably damaging | 17.2 | Pathogenic |
| R100Q | c.299G>A | 90132 | - | 1.0 Probably damaging | 14.2 | Pathogenic |
| A111P | c.331G>C | 90170 | - | 0.999 Probably damaging | 14.4 | Pathogenic |
| H112D | c.334C>G | 628998 | - | 1.0 Probably damaging | 4.31 | Neutral |
| L135V | c.403C>G | - | - | 0.693 Possibly damaging | 4.31 | Neutral |
| E234G | c.701A>G | - | - | 0.005 Benign | 2.19 | Neutral |
| G244V | c.731G>T | 90340 | - | 0.512 Possibly damaging | 6.84 | Pathogenic |
| K286Q | c.856A>C | - | - | 0.986 Probably damaging | 1.61 | Pathogenic |
| N338S | c.1013A>G | 89605 | - | 0.140 Benign | 2.78 | Neutral |
| G454R | c.1360G>C | 89705 | 1.11E-2 | 0.000 Benign | 4.47 | Neutral |
| R474P | c.1421G>C | - | 0.489 | 0.007 Benign | - | Neutral |
| R474Q | c.1421G>A | 89737 | - | 0.022 Benign | - | Neutral |
| V506A | c.1517T>C | 89757 | - | 0.980 Probably damaging | 5.56 | Neutral |
| Q542L | c.1625A>T | 89811 | - | 1.0 Probably damaging | 24.2 | Pathogenic |

| | | | | | | |
|---|---|---|---|---|---|---|
| S627T | c.1879T>A | - | - | 0.890 Possibly damaging | 6.32 | Pathogenic |
| D748A | c.2243A>C | - | - | 0.951 Probably damaging | 4.51 | Neutral |
| *Pathogenic* | | | | | | |
| I19F | c.55A>T | 90274 | 346 | 0.992 Probably damaging | 9.59 | Pathogenic |
| D41H | c.121G>C | 89682 | 3.99E+4 | 1.0 Probably damaging | 20.3 | Pathogenic |
| S44F | c.131C>T | 17079 | 2.46E+4 | 0.953 Probably damaging | 16.7 | Pathogenic |
| G67R | c.199G>A | 89992 | 4.27E+4 | 1.0 Probably damaging | 36.5 | Pathogenic |
| R100P | c.299G>C | 90133 | 1,450 | 1.0 Probably damaging | 40.3 | Pathogenic |
| I107R | c.320T>G | 90167 | 210 | 0.995 Probably damaging | 34.8 | Pathogenic |
| T117M | c.350C>T | 17094 | 1.33E+18 | 0.997 Probably damaging | 21.1 | Pathogenic |
| S247P | c.739T>C | 90342 | 105 | 0.976 Probably damaging | 7.88 | Pathogenic |
| R265S[e] | c.793C>A | 90380 | 37.2 | 0.999 Probably damaging | 13.6 | Pathogenic |
| I276R | c.827T>G | 520540 | 319 | 0.964 Probably damaging | 32.2 | Pathogenic |
| R687W | c.2059C>T | 90014 | 1.74E+4 | 0.963 Probably damaging | 3.87 | Pathogenic |

[a] Odds of Pathogenicity calculated based on known clinical and personal data from patients harboring variant. Any prior functional data was excluded in the calculation. Input data for each variant can be found at http://insight-database.org/classifications/. I291V and H718Y were classified as benign due to frequency of variant in the population.

[b] PolyPhen-2 prediction scores were calculated from http://genetics.bwh.harvard.edu/pph2/[7]. Scores range between 0-1. Scores close to 0 are predicted to be benign while scores close to 1 indicate damaging impact on protein function.

[c] MAPP-MMR prediction scores obtained from http://mappmmr.blueankh.com/[8]. Higher score indicates greater predicted deleterious impact on protein function. Scores > 3.5 are considered either VUS or deleterious in nature. There is no available MAPP-MMR score for the R474P and R474Q variants.

[d] PON-MMR2 predictions obtained from http://structure.bmc.lu.se/PON-MMR2/[9]

[e] R265S is a pathogenic variant when prior functional data are used in the classification; without the use of these data, the variant would be classified as a likely pathogenic variant

**Supplementary Table 3 Primers used to genotype the *MLH1* variant carrying cell lines**

| Cell line | Forward primer | Reverse primer |
|---|---|---|
| *Benign* | | |
| D132H | CCTTGGGATTAGTATCTATCTCTC | CCATGCCACAAAAGCCAATAGTC |
| V213M | AGTTTGCTGGTGGAGATAAGG | GAACACATGATTCACGCCAC |
| I219V | TAGTTTGCTGGTGGAGATAAGG | CAAGCCTGTGTATTTGACTAAAGC |
| E268G | GGAAAGTGGCGACAGGTAAA | TCAAAGAGGAGAGCCTGATAGA |
| S406N | ACCAGATGGTTCGTACAGATTC | GGAAGTAGGTCCTCTCTTCTCT |
| E578G | TGTCTGTCGTATCTTCTCTGTTG | GCTGTCACACCTCATCAATTTC |
| K618A | CTTGCTTCTTCCTAGGAGCC | GCTGTCACACCTCATCAATTTC |
| Q689R | CAGGTGAATTGGGACGAAGA | GAGGTCACTCTGGAGGTATGT |
| S692P | CAGGTGAATTGGGACGAAGA | CCTTGTTGGGTACAAAGAAATCC |
| V716M | CTAGCCCACAAGATCCACTTC | ATGTGTGAGCGCAAGGCTTTA |
| H718Y | TAACACCAAGTCTTTCCAGACC | CACATCCCACAGTGCATAAATAAC |
| *VUS* | | |
| R9W | CGCAAGCGCATATCCTTCTA | GAGATGATTGAGAACTGGTACGG |
| C39R | GCACTTCCGTTGAGCATCTA | GGGAGAGCGGTAAAGAAACA |
| A42V | CAGTGTATGAGCCTGTAAGACAA | GGCCTCCCTCTTTAACAATCA |
| D63N | GTATGAGCCTGTAAGACAAAGGA | GAACAGTGCCCAGCAAATAATAG |
| N64S | GTATGAGCCTGTAAGACAAAGGA | GAACAGTGCCCAGCAAATAATAG |
| L73P | ATGGGAATTCAAAGAGATTTGG | ATGTCATCACAGGAGGATATTT |
| R100Q | ACTAGTAAACTGCAGTCCTTTGA | TGGGTACTTCCATCAAGAAACA |
| A111P | GGTGACCCAGCAGTGAGTTT | ACTGGTGTTGAGACAGGATTAC |
| H112D | GGTGACCCAGCAGTGAGTTT | ACTGGTGTTGAGACAGGATTAC |
| L135V | CCCTTGGGATTAGTATCTATCTCTCTA | GGATATCTTGGGACCTCCATTAAC |
| E234G | TGGGAAGGAACCTTGTGTTT | CAAACTTTGCCATGAGGTTTCT |
| G244V | TGGGAAGGAACCTTGTGTTT | CTGAGCACAGACTTAGGACAAA |
| K286Q | TGTGAATGTACACCTGTGACC | CAAAGAGGAGAGCCTGATAGAAC |
| N338S | CCCAGAATGTGGATGTTAATGTG | AGGCAAAGTGAGGAAGTGAG |
| G454R | CAGGCCATTGTCACAGAGG | GCAGAGAGAAGATGCAAGTGA |
| R474P | CTGTAGAACCAGCACAGAGAAG | CAACATGACTGCTTTCTCCATTT |
| R474Q | CTGTAGAACCAGCACAGAGAAG | CAACATGACTGCTTTCTCCATTT |
| V506A | CAGAAAGAGACATCGGGAAGAT | GTGGGTTAGTAAAGGAAGAGGAG |
| Q542L | TCCGGGAGATGTTGCATAAC | GGAAATGGTCGAACTTGGATTTG |
| S627T | GGTTTCTCACCTGCCATTCT | TCCCAAAGTGCTGGGATTAC |
| D748A | ACACCAGTGTATGTTGGGATG | AGGAATACTATCAGAAGGCAAGTATAA |
| *Pathogenic* | | |
| I19F | CGCAAGCGCATATCCTTCTA | CCGTACCAGTTCTCAATCATCTC |
| D41H | AGAAATGATGGTTGCTCTGCCC | TACTTACCCTGATCCCGGTGC |
| S44F | TGAGCCTGTAAGACAAAGGAAA | CCATGAAGCGCACAAACATC |
| G67R | TGCCAGTTTAGATGCAAAATCC | GAACAGTGCCCAGCAAATAATAG |
| R100P | ACTAGTAAACTGCAGTCCTTTGA | TGGGTACTTCCATCAAGAAACA |
| I107R | GGTGACCCAGCAGTGAGTTT | ACTGGTGTTGAGACAGGATTAC |
| T117M | CATCAAAGCAAGTGAGCAAAGT | CGTACTCAAGATCTCTGCCAAA |
| S247P | TGGGAAGGAACCTTGTGTTT | ATTCCCTGTGGGTGTTTCCT |
| R265S | GGAAAGTGGCGACAGGTAAA | TCAAAGAGGAGAGCCTGATAGA |
| I276R | CAAAGGGAAGTGAGGAGGGA | TGAGGAGTTTGGTGCTACATTAC |
| R687W | CAGGTGAATTGGGACGAAGA | AGGTCACTCTGGAGGTATGT |

### Benign variants

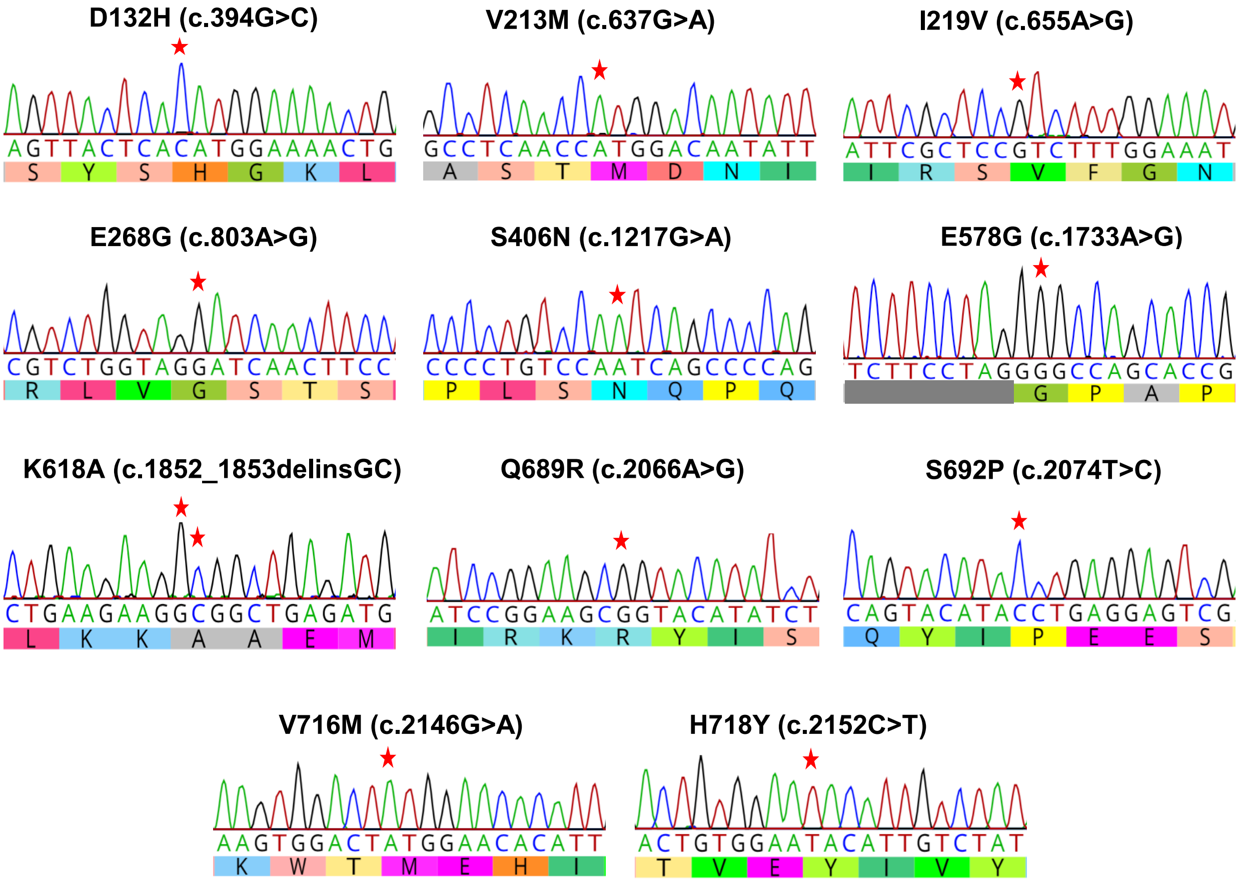

### Variants of uncertain significance

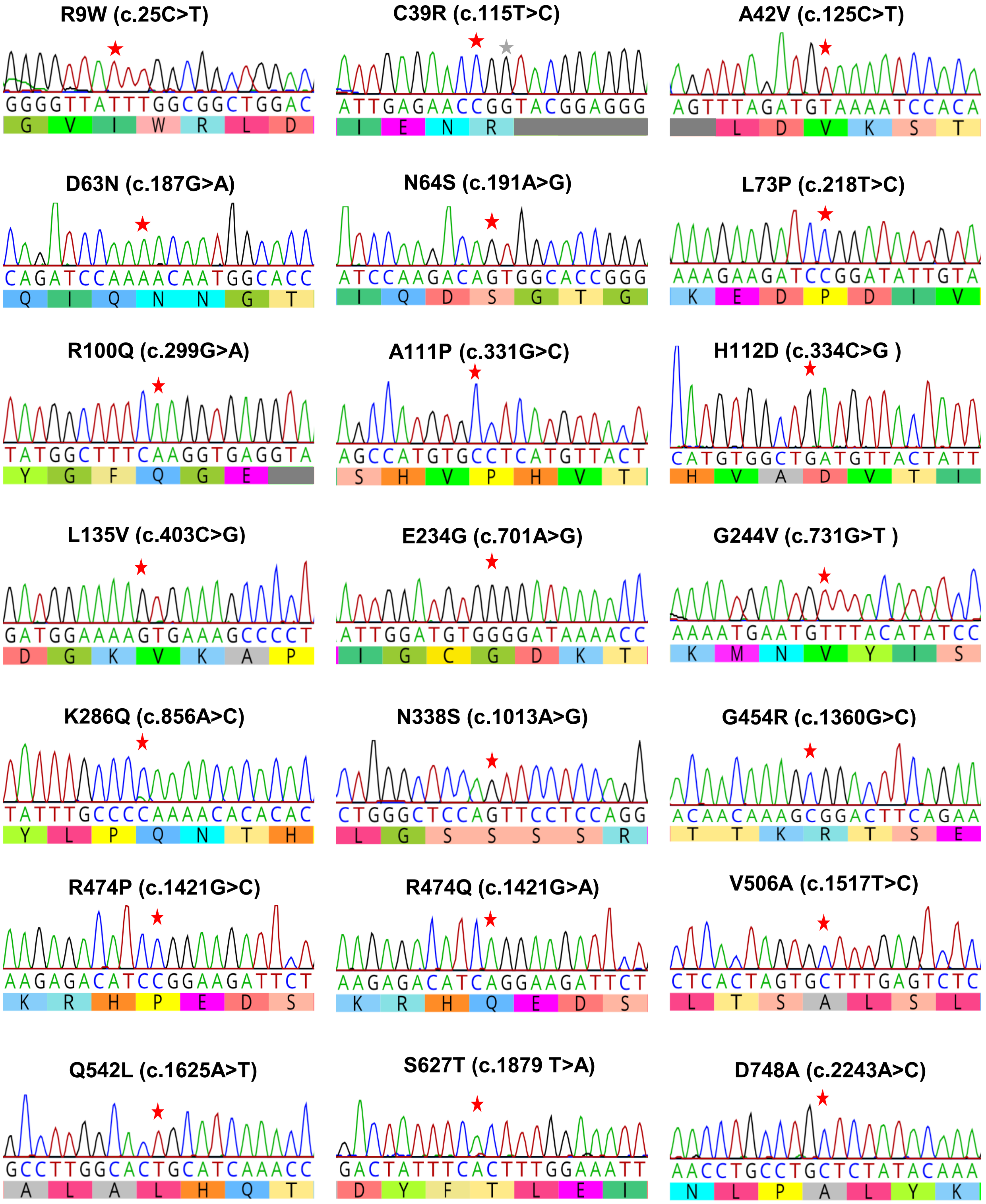

### Pathogenic variants

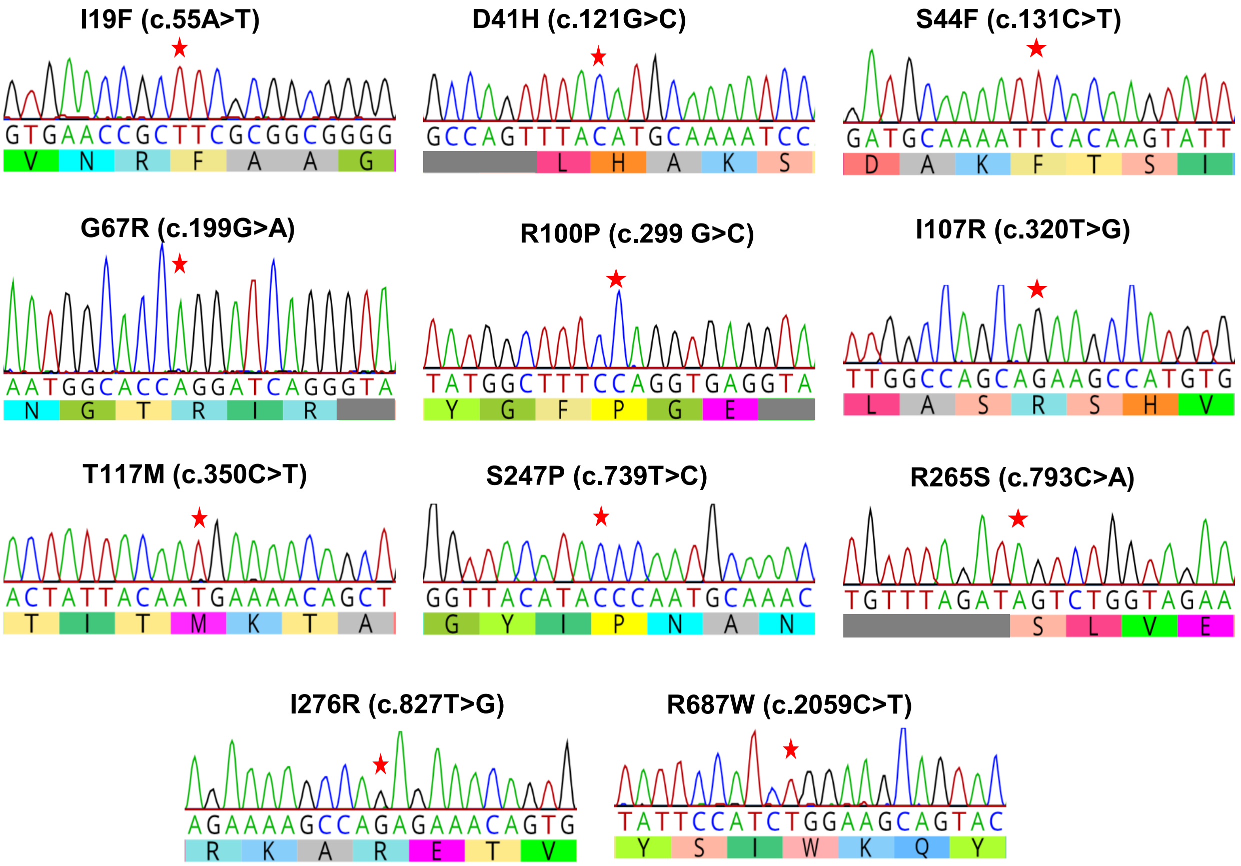


**Fig. S1. Sanger sequencing results of the targeted region in all homozygous *MLH1* variant expressing cell lines.** Images display DNA chromatograms for each generated cell line showing homozygous insertion of the codon-altering variants (red stars).

**Supplementary Table 4 *MLH1* variant targeting efficiencies**

| Cell line | Method | Nuclease | No. of homozygous clones | No. of clones screened | Distance from PAM | Targeting Efficiency (%) |
|---|---|---|---|---|---|---|
| *Benign* | | | | | | |
| D132H | Plasmid | Cas9 | 5 | 46 | 2 | 11 |
| V213M | RNP | Cas9 | 2 | 56 | 1 | 4 |
| I219V | RNP | Cas12a | 6 | 32 | 5 | 19 |
| E268G | RNP | Cas12a | 3 | 21 | 14 | 14 |
| S406N | Plasmid | Cas9 | 1 | 48 | 1 | 2 |
| E578G | RNP | Cas9 | 2 | 40 | 5 | 5 |
| K618A | RNP | Cas12a | 5 | 48 | 9 | 10 |
| Q689R | Plasmid | Cas9 | 1 | 20 | 6 | 5 |
| S692P | RNP | Cas9 | 6 | 44 | 4 | 14 |
| V716M | RNP | Cas9 | 1 | 32 | 1 | 3 |
| H718Y | RNP | Cas12a | 5 | 48 | 11 | 10 |
| *VUS* | | | | | | |
| R9W | RNP | Cas9 | 1 | 2 | 3 | 50 |
| C39R | RNP | Cas9 | 4 | 40 | 5 | 10 |
| A42V | RNP | Cas12a | 12 | 32 | 5 | 38 |
| D63N | RNP | Cas9 | 2 | 56 | 5 | 4 |
| N64S | RNP | Cas9 | 5 | 48 | 1 | 10 |
| L73P | RNP | Cas12a | 12 | 64 | 14 | 19 |
| R100Q | RNP | Cas12a | 4 | 56 | 1 | 7 |
| A111P | Plasmid | Cas9 | 2 | 44 | 5 | 5 |
| H112D | RNP | Cas12a | 8 | 48 | 17 | 17 |
| L135V | RNP | Cas12a | 7 | 64 | 2 | 11 |
| E234G | RNP | Cas12a | 3 | 64 | 4 | 5 |
| G244V | RNP | Cas12a | 6 | 56 | 15 | 11 |
| K286Q | RNP | Cas12a | 1 | 48 | 4 | 2 |
| N338S | Plasmid | Cas9 | 2 | 14 | 1 | 14 |
| G454R | RNP | Cas9 | 1 | 40 | 0 | 3 |
| R474P | RNP | Cas12a | 3 | 48 | 17 | 6 |
| R474Q | RNP | Cas12a | 3 | 24 | 17 | 13 |
| V506A | RNP | Cas12a | 5 | 48 | 17 | 10 |
| Q542L | Plasmid | Cas9 | 4 | 48 | 7 | 8 |
| S627T | RNP | Cas12a | 0 | 40 | 1 | 0[a] |
| D748A | RNP | Cas12a | 6 | 48 | 7 | 13 |
| *Pathogenic* | | | | | | |
| I19F | Plasmid | Cas9 | 7 | 32 | 4 | 22 |
| D41H | RNP | Cas12a | 5 | 56 | 1 | 9 |
| S44F | RNP | Cas12a | 5 | 18 | 11 | 28 |
| G67R | RNP | Cas9 | 2 | 48 | 5 | 4 |
| R100P | RNP | Cas12a | 7 | 31 | 1 | 23 |
| I107R | Plasmid | Cas9 | 1 | 32 | 4 | 3 |
| T117M | RNP | Cas12a | 2 | 49 | 1 | 4 |
| S247P | RNP | Cas12a | 3 | 27 | 7 | 11 |
| R265S | RNP | Cas12a | 9 | 63 | 5 | 14 |
| I276R | RNP | Cas12a | 1 | 40 | 2 | 3 |
| R687W | RNP | Cas9 | 2 | 47 | 3 | 4 |

[a] Only heterozygous clones were initially obtained that were re-targeted to generate homozygous clones

| | | | | | | |
|---|---|---|---|---|---|---|
| R687W | RNP | Cas9 | 2 | 47 | 3 | 4 |

[a] Only heterozygous clones were initially obtained that were re-targeted to generate homozygous clones

**Supplementary Table 5 Examination of Top Five Candidate Loci for Off-target Cleavage by CRISPR Nuclease**

| Cell Line | I | II | III | IV | V |
|---|---|---|---|---|---|
| *Benign* | | | | | |
| D132H | | | | | |
| V213M | | | | | |
| I219V | | | | | |
| E268G | | | | | |
| S406N | | | | | |
| E578G | | | | | |
| K618A | | | | | |
| Q689R | | | | | |
| S692P | | | | | |
| V716M | | | | | |
| H718Y | | | | | |
| *VUS* | | | | | |
| R9W | | | | | |
| C39R | | | | | |
| A42V | | | | | |
| D63N | | | | | |
| N64S | | | | | |
| L73P | | | | | |
| R100Q | | | | | |
| A111P | | | | | |
| H112D | | | | | |
| L135V | | | | | |
| E234G | | | | | |
| G244V | | | | | |
| K286Q | | | | | |
| N338S | | | | | |
| G454R | | | | | |
| R474P | | | | | |
| R474Q | | | | | |
| V506A | | | | | |
| Q542L | | | | | |
| S627T | | | | | |
| D748A | | | | | |
| *Pathogenic* | | | | | |
| I19F | | | | | |
| D41H | | | | | |
| S44F | | | | | |
| G67R | | | | | |
| R100P | | | | | |
| I107R | | | | | |
| T117M | | | | | |
| S247P | | | | | |
| R265S | | | | | |
| I276R | | | | | |
| R687W | | | | | |
| MLH1 KO | | | | | |

Green – Sequence at candidate off-target site matches expected WT sequence
Black – K618A sgRNA had less than five candidate off-target sites

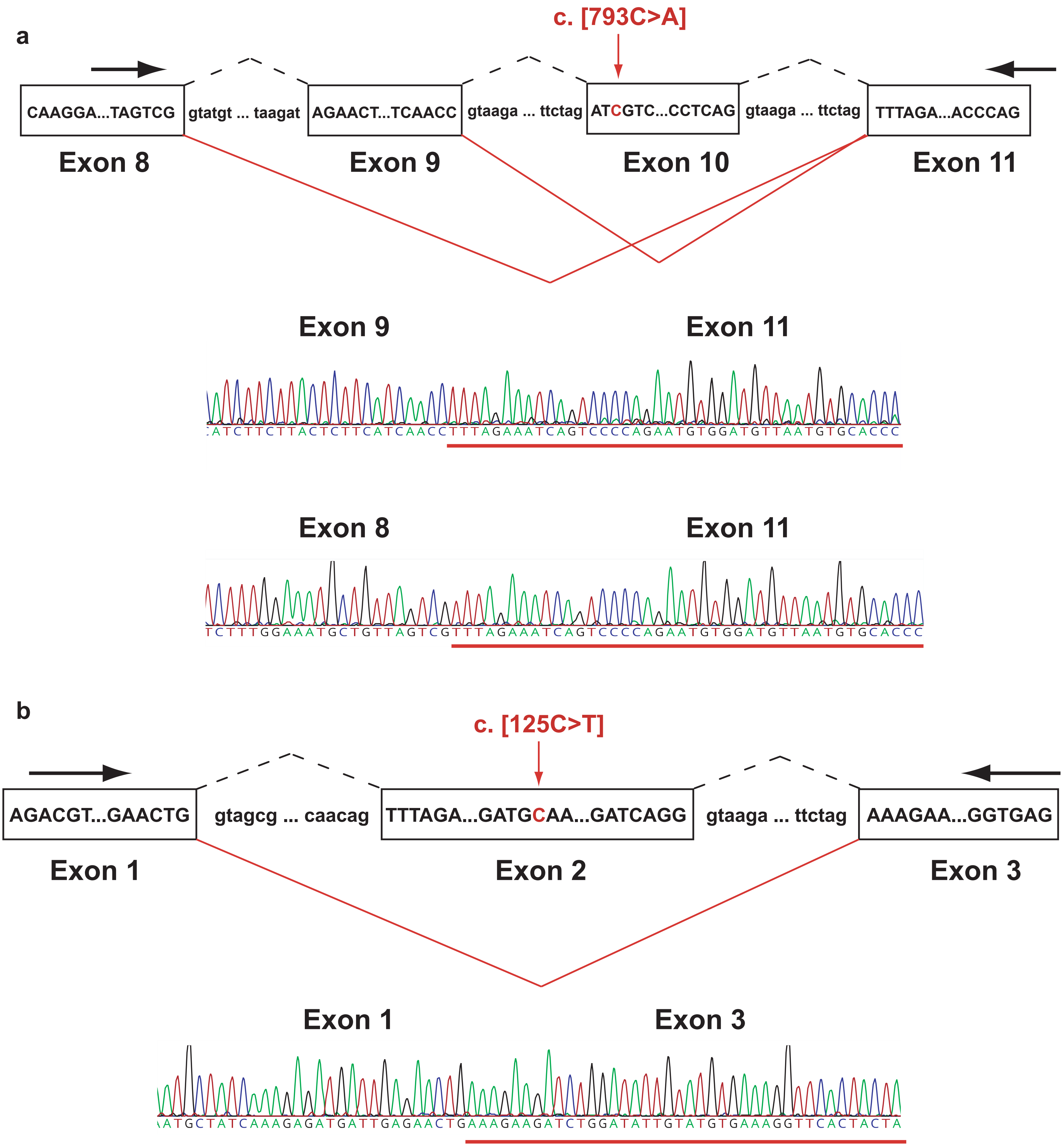


**Fig. S2. Aberrant exon splicing in R265S and A42V cell lines.** Schematic of exons that are aberrantly spliced due to the R265S (**a**) and A42V (**b**) variants. The nucleotide variant is shown in red. The black dotted lines indicate splicing as observed in canonical WT transcripts. The solid red lines indicate variant induced exon exclusion. The Sanger sequencing chromatograms are also shown for each aberrant splice product. Arrows indicate the position of PCR primers used in sequencing the cDNA splice products.

**a**

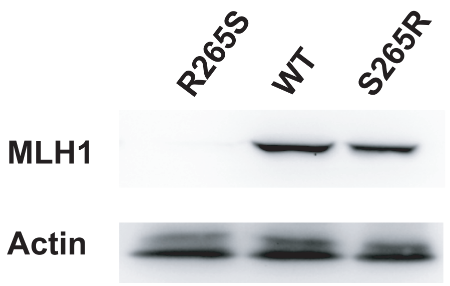


**b**

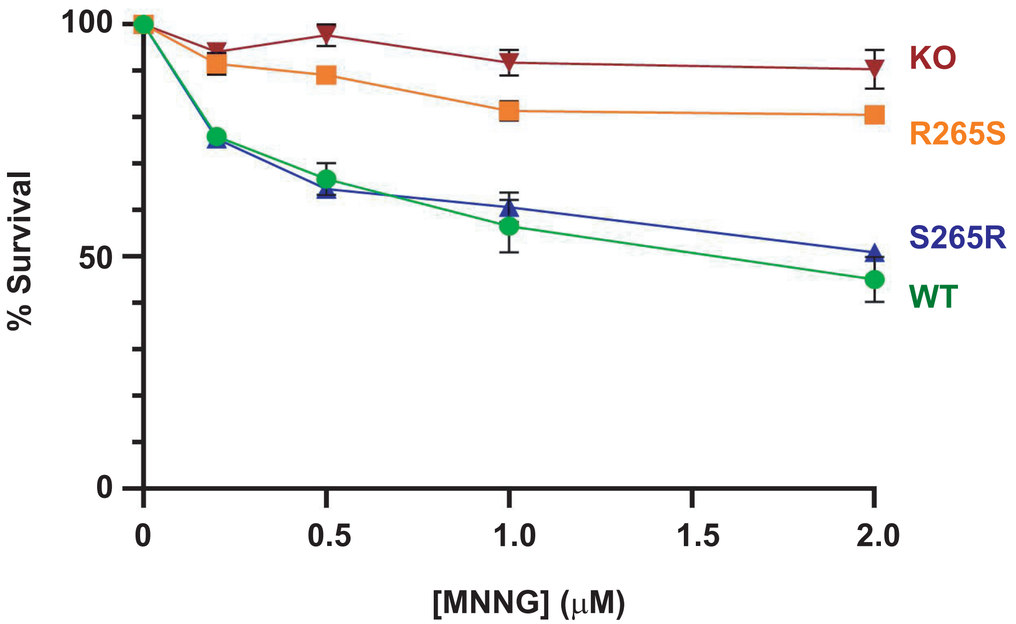


**Fig. S3. Revertent S265R variant restores normal protein levels and response to DNA damage. (a)** Western blot with an anti-MLH1 antibody and anti-actin antibody as loading control showing that correction of the R265S variant back to the WT sequence leads to restoration of normal MLH1 protein levels. **(b)** MNNG survival assay showing that revertent S265R variant restores the DNA damage response following a 24 hr treatment at various concentrations of MNNG. n=3.

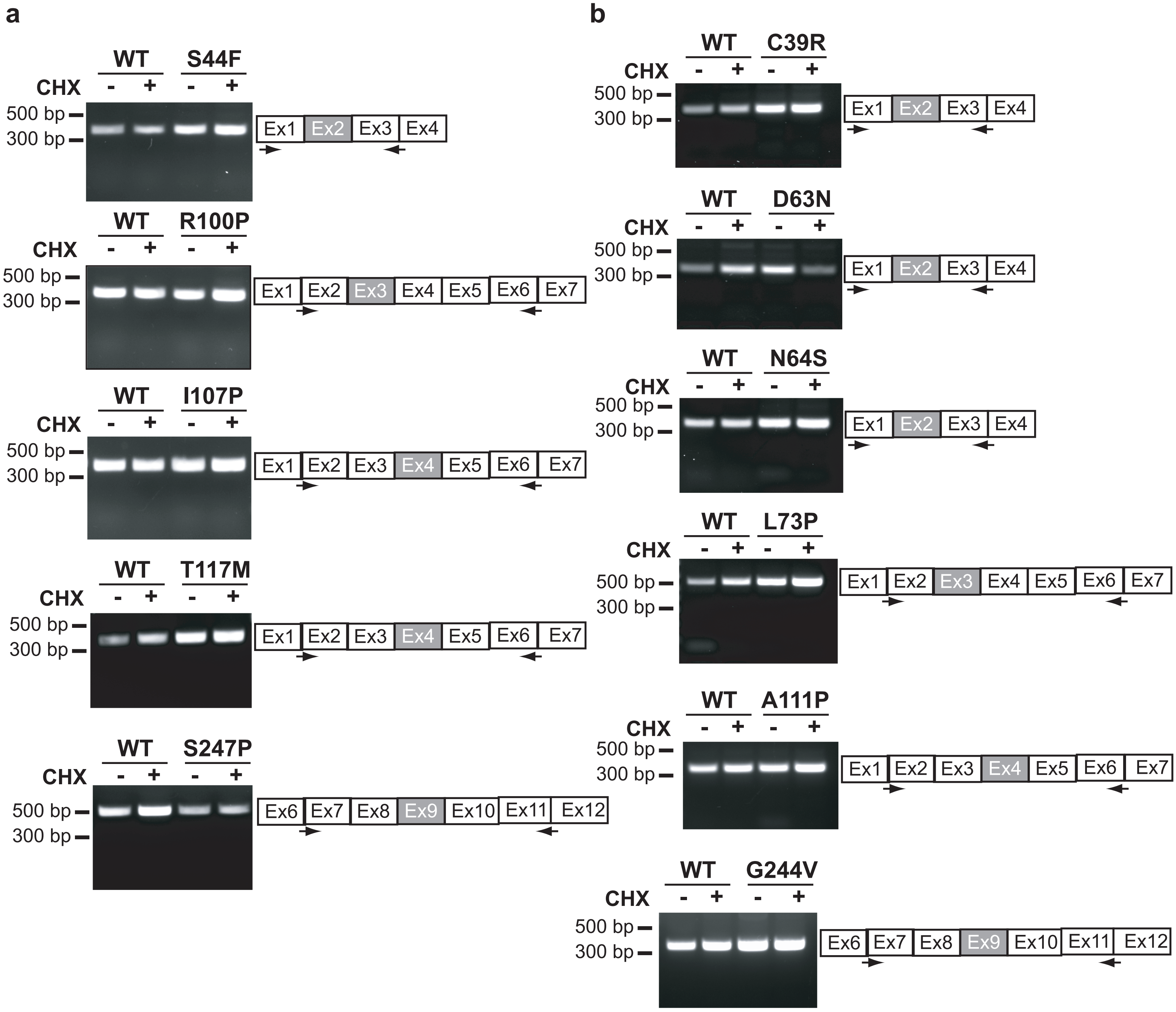


**Fig. S4. Exon exclusion assay results for *MLH1* variants.** Reverse transcribed total RNA from pathogenic variant cells **(a)** and VUS cells **(b)** with reduced MLH1 protein levels was PCR amplified demonstrating that the indicated variants caused no aberrant splicing. CHX, Cycloheximide.

**Supplementary Table 6 Primers used for exon exclusion assay**

| Cell line | Forward primer | Reverse primer |
|---|---|---|
| *VUS* | | |
| C39R | GACGTTTCCTTGGCTCTTCTG | CTTATGCTGGCCAAAGCCTC |
| A42V | GACGTTTCCTTGGCTCTTCTG | CTTATGCTGGCCAAAGCCTC |
| D63N | GACGTTTCCTTGGCTCTTCTG | CTTATGCTGGCCAAAGCCTC |
| N64S | GACGTTTCCTTGGCTCTTCTG | CTTATGCTGGCCAAAGCCTC |
| L73P | GATCCAAGACAATGGCACCGG | GTGTACTGAATACCTGCCAAC |
| A111P | CGGGATCAGGAAAGAAGATCT | GTGTACTGAATACCTGCCAAC |
| G244V | GAAGTTGTTGGCAGGTATTCAG | GTAGCAAAGTCTGGGTGAAGTAC |
| *Pathogenic* | | |
| S44F | GACGTTTCCTTGGCTCTTCTG | CTTATGCTGGCCAAAGCCTC |
| R100P | GATCCAAGACAATGGCACCGG | GTGTACTGAATACCTGCCAAC |
| I107R | CGGGATCAGGAAAGAAGATCT | GTGTACTGAATACCTGCCAAC |
| T117M | CGGGATCAGGAAAGAAGATCT | GTGTACTGAATACCTGCCAAC |
| S247P | GAAGTTGTTGGCAGGTATTCAG | GTAGCAAAGTCTGGGTGAAGTAC |
| R265S | GCAGGTATTCAGTACACAATGC | CTGGGTGAAGTACATCCTGG |

**Supplementary Table 7 Primers used in the MSI assay**

| Marker | Forward Primer | Reverse Primer | Fluoro-phore |
|---|---|---|---|
| NR-27 | AACCATGCTTGCAAACCACT | **G**CGATAATACTAGCAATGACC | 6-FAM |
| NR-21 | CTGGTCACTCGCGTTTACAA | GAGTCGCTGGCACAGTTCTA | HEX |
| NR-22 | CGAGGCTTGTCAAGGACATAA | GCCATCCAGTTTTGTTCTTACA | 6-FAM |
| BAT-25 | CTTTCCTCGCCTCCAAGAATG | CCACACTTCAAAATGACATTCTGC | HEX |
| BAT-26 | CTGCGGTAATCAAGTTTT | **G**AACCATTCAACATTTTTAACCC | 6-FAM |

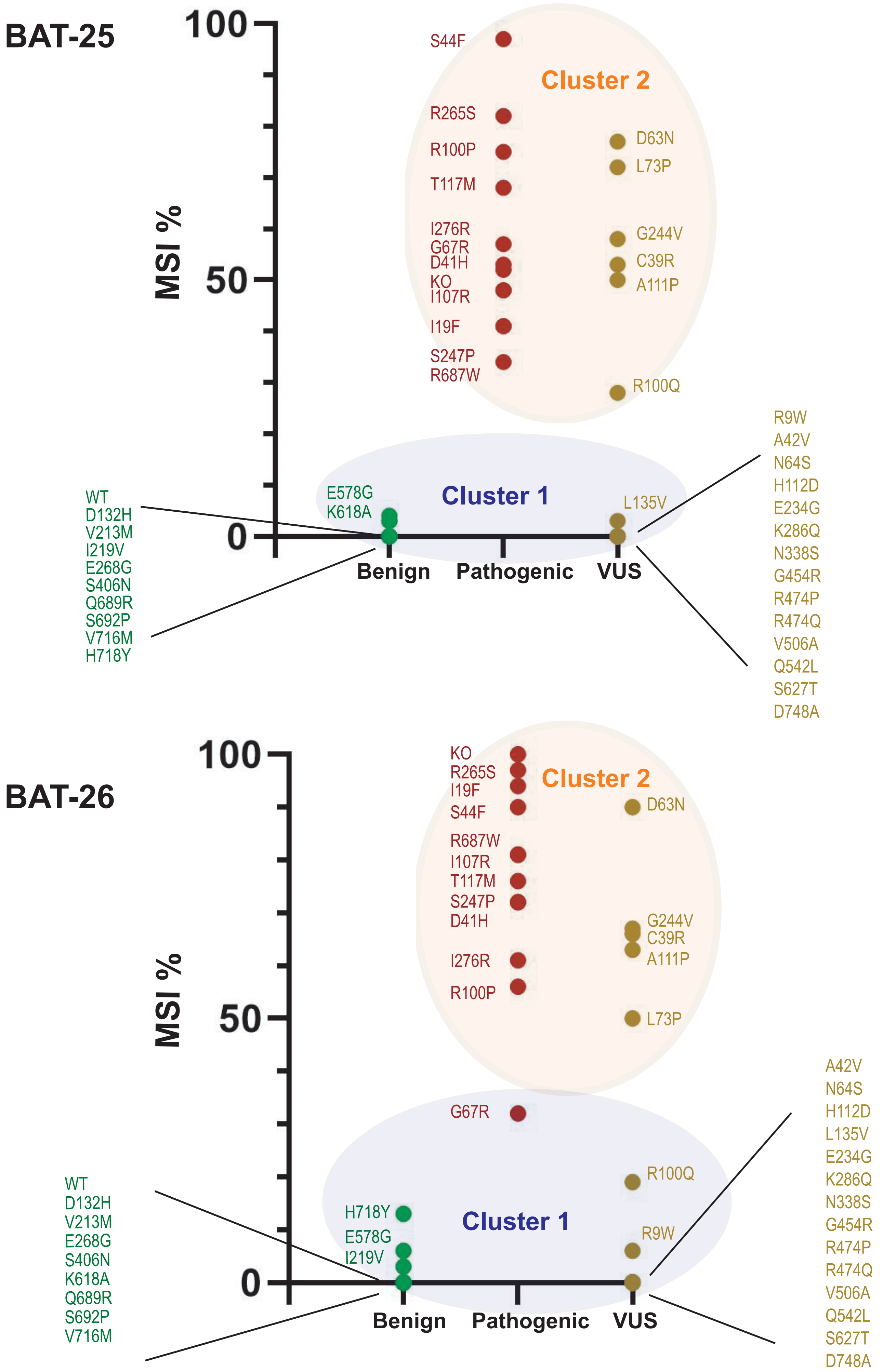
BAT-25
100
50
0
MSI %
Cluster 2
S44F
R265S
R100P
T117M
I276R
G67R
D41H
KO
I107R
I19F
S247P
R687W
D63N
L73P
G244V
C39R
A111P
R100Q
Cluster 1
E578G
K618A
L135V
WT
D132H
V213M
I219V
E268G
S406N
Q689R
S692P
V716M
H718Y
R9W
A42V
N64S
H112D
E234G
K286Q
N338S
G454R
R474P
R474Q
V506A
Q542L
S627T
D748A
Benign
Pathogenic
VUS
BAT-26
100
50
0
MSI %
Cluster 2
KO
R265S
I19F
S44F
R687W
I107R
T117M
S247P
D41H
I276R
R100P
D63N
G244V
C39R
A111P
L73P
G67R
R100Q
Cluster 1
H718Y
E578G
I219V
R9W
WT
D132H
V213M
E268G
S406N
K618A
Q689R
S692P
V716M
A42V
N64S
H112D
L135V
E234G
K286Q
N338S
G454R
R474P
R474Q
V506A
Q542L
S627T
D748A
Benign
Pathogenic
VUS

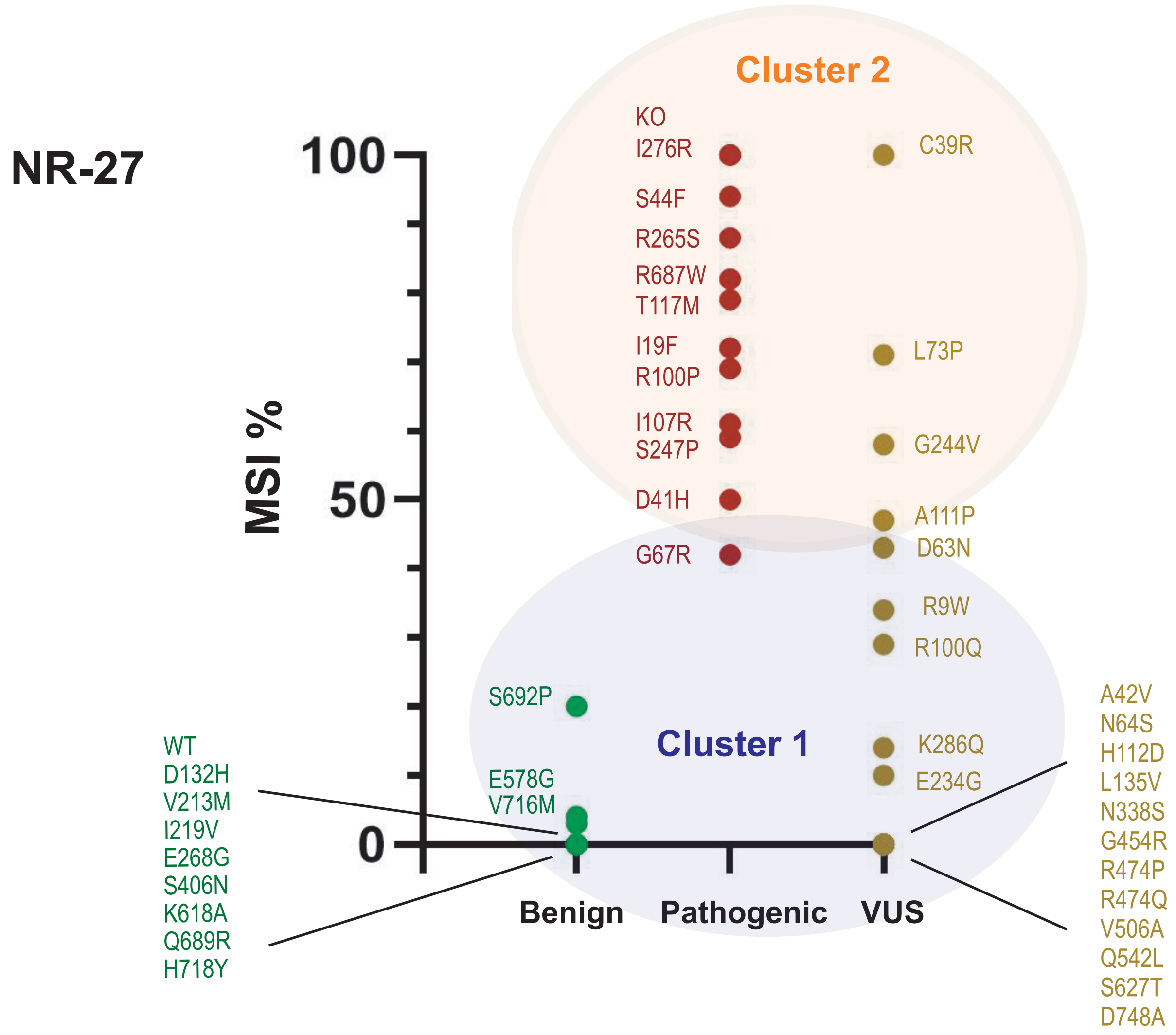

NR-27
Cluster 2
Cluster 1
MSI %
100
50
0
Benign
Pathogenic
VUS
KO
I276R
S44F
R265S
R687W
T117M
I19F
R100P
I107R
S247P
D41H
G67R
C39R
L73P
G244V
A111P
D63N
R9W
R100Q
K286Q
E234G
S692P
E578G
V716M
WT
D132H
V213M
I219V
E268G
S406N
K618A
Q689R
H718Y
A42V
N64S
H112D
L135V
N338S
G454R
R474P
R474Q
V506A
Q542L
S627T
D748A

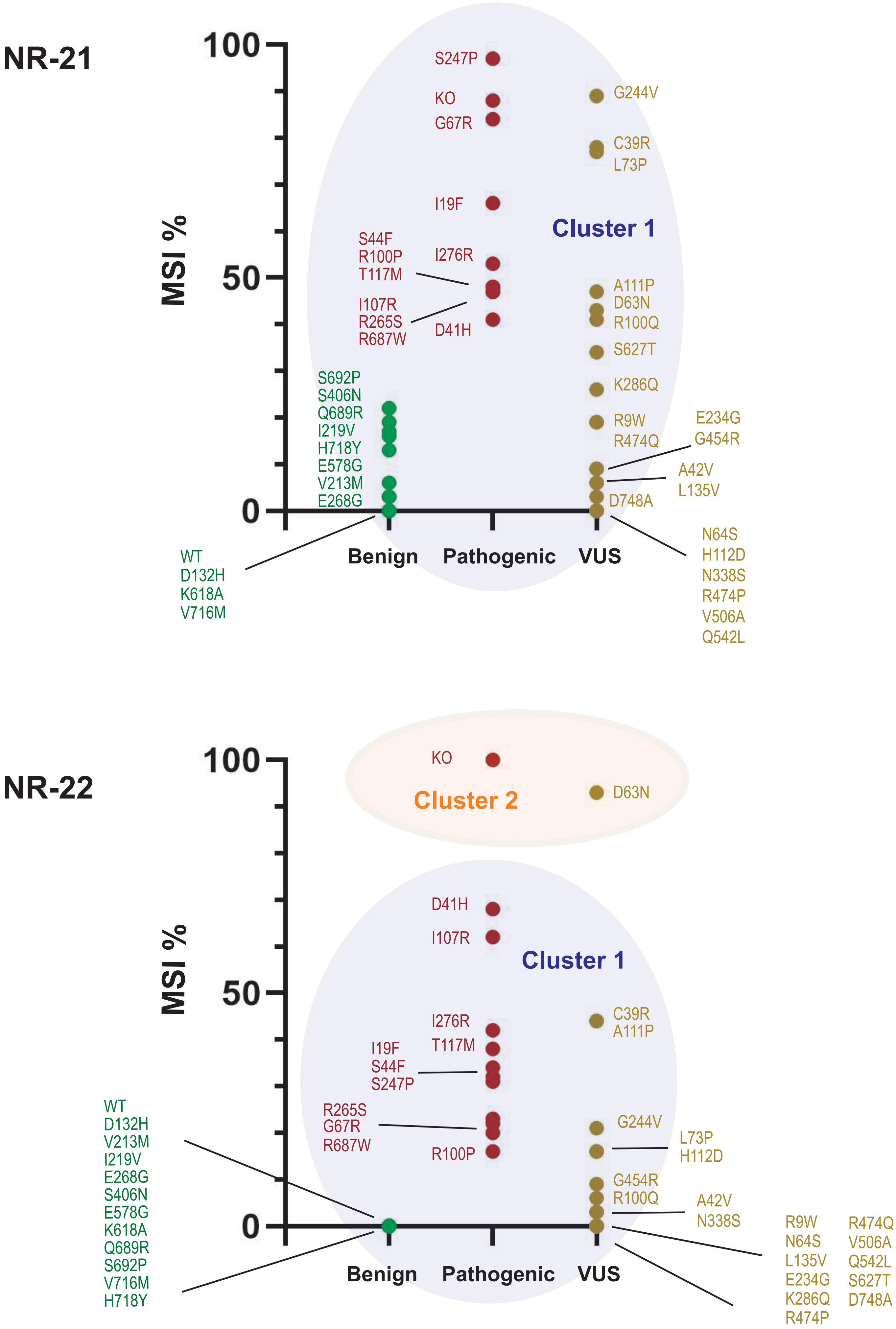

NR-21
MSI %
100
50
0
Benign
Pathogenic
VUS
Cluster 1
S247P
KO
G67R
I19F
I276R
S44F
R100P
T117M
I107R
R265S
R687W
D41H
G244V
C39R
L73P
A111P
D63N
R100Q
S627T
K286Q
R9W
R474Q
E234G
G454R
A42V
L135V
D748A
N64S
H112D
N338S
R474P
V506A
Q542L
S692P
S406N
Q689R
I219V
H718Y
E578G
V213M
E268G
WT
D132H
K618A
V716M
NR-22
MSI %
100
50
0
Benign
Pathogenic
VUS
KO
Cluster 2
D63N
Cluster 1
D41H
I107R
I276R
T117M
I19F
S44F
S247P
R265S
G67R
R687W
R100P
C39R
A111P
G244V
L73P
H112D
G454R
R100Q
A42V
N338S
R9W
N64S
L135V
E234G
K286Q
R474P
R474Q
V506A
Q542L
S627T
D748A
WT
D132H
V213M
I219V
E268G
S406N
E578G
K618A
Q689R
S692P
V716M
H718Y

**Fig. S5. Statistical clustering of microsatellite instability (MSI) results.** Graphs demonstrating percentage of unstable clones for the pathogenic and benign variants and VUS from five MSI markers: BAT-25, BAT-26, NR-27, NR-21 and NR-22. Statistical clustering was performed to segregate the results into two separate clusters (blue and orange circles). For three markers (BAT-25, BAT-26 and NR-27), the pathogenic and benign controls mostly segregated into two separate clusters (with two discordant results), while these controls failed to segregate cleanly for the NR-21 and NR-22 markers.
